# Benchmark of biomarker identification and prognostic modeling methods on diverse censored data

**DOI:** 10.64898/2026.03.29.715113

**Authors:** Wesley Fletcher, Samiran Sinha

## Abstract

The practices of identifying biomarkers and developing prognostic models using genomic data has become increasingly prevalent. Such data often features characteristics that make these practices difficult, namely high dimensionality, correlations between predictors, and sparsity. Many modern methods have been developed to address these problematic characteristics while performing feature selection and prognostic modeling, but a large-scale comparison of their performances in these tasks on diverse right-censored time to event data (aka survival time data) is much needed. We have compiled many existing methods, including some machine learning methods, several which have performed well in previous benchmarks, primarily for comparison in regards to variable selection capability, and secondarily for survival time prediction on many synthetic datasets with varying levels of sparsity, correlation between predictors, and signal strength of informative predictors. For illustration, we have also performed multiple analyses on a publicly available and widely used cancer cohort from The Cancer Genome Atlas using these methods. We evaluated the methods through extensive simulation studies in terms of the false discovery rate, F1-score, concordance index, Brier score, root mean square error, and computation time. Of the methods compared, CoxBoost and the Adaptive LASSO performed well in all metrics, and the LASSO and elastic net excelled when evaluating concordance index and F1-score. The Benjamini-Hoschberg and q-value procedures showed volatile performances in controlling the false discovery rate. Some methods’ performances were greatly affected by differences in the data characteristics. With our extensive numerical study, we have identified the best performing methods for a plethora of data characteristics using informative metrics. This will help cancer researchers in choosing the best approach for their needs when working with genomic data.

## Introduction

In cancer genomics studies, some common goals are the identification of biomarkers, which can be used for early diagnosis and improved prognosis (e.g. predicted time until death) of individuals diagnosed with the disease. Studies that pursue these goals frequently collect time-to-event data containing gene expression data (the covariates or features) and the survival time (the response) of each observational unit. A widely accepted statistical model to relate these covariates with the response is the Cox proportional hazards model. Common characteristics of such time-to-event data are:

- right-censoring, where if an observational unit stops being observed before the time of failure then it is called “censored” or “right-censored.”
- high dimensionality, where the number of covariates exceeds the number of observational units (i.e. “high-*p*, low-*n*”).
- correlation between covariates.
- sparse covariates (i.e. few of the observed covariates are useful and contribute to differences in survival time).

These characteristics can make working with such data a very difficult task. Various statistical and machine learning methods have been developed to perform feature selection (identifying a handful of features out of a large number of features) while fitting the Cox proportional hazards model to the time-to-event. Methods that can effectively accomplish these tasks may be instrumental in identifying biomarkers and performing prognosis. As such, we seek to evaluate the performances of many of these methods on synthetic data featuring the characteristics listed above in hopes of identifying the most suitable ones for practitioners to employ.

Typically, the data is presented as *n* independent observations of (*Y*, Δ, *X*), where the response variable *Y* = min(*T, C*) denotes the minimum of the time-to-event *T* and the censoring time *C*, indicator Δ = *I*(*T* ≤ *C*) takes on zero or one for right-censored and uncensored subject respectively, and *X* denotes a *p* vector of covariates for a random subject. We add the suffix *i* to every random variable to identify data from the *i*th observational unit and *i* = 1, …, *n* as we presumed to have data from *n* observational units. In our context, *p* is larger than *n*.

Under the Cox proportional hazards model [1], the hazard of the time-to-event for observation *i* at time *t* is

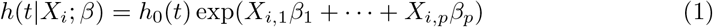

where *h*_0_(*t*) ≥ 0 is the baseline hazard at time *t*, and *β* = (*β*_1_, ···, *β*_*p*_)^⊤^ is a vector of regression parameters, which relates the length *p* covariate *X*_*i*_ = (*X*_*i*,1_, ···, *X*_*i,p*_)^⊤^ with the hazards of the event. Note that a zero value of a component of the *β*-vector implies the corresponding covariate is not related with the hazards. Inversely, the covariates corresponding to the non-zero components of *β* are the important or regulatory genes for the hazards, and can be considered prognostic markers. The main goal is identification of the non-zero components of the *β*-vector from the observed data where the sample size is smaller than the number of covariates/features. Moreover, a positive (negative) value of a regression parameter implies that a larger (smaller) value of the feature is associated with a higher hazard.

Although past benchmarks have been performed, our novelty lies in 1) encompassing a greater variety of methods for comparison, 2) the diversity of our synthetic datasets, and 3) the emphasis on simultaneously evaluating feature selection and predictive capabilities. A brief summary of pertinent literature is given below.

Saeys et al. [2] gave an extensive taxonomy and discussion of filter, wrapper, and embedded feature selection techniques. Hira et al. [3] examined various examples of filter, wrapper, and embedded techniques and analyze their specific strengths and weaknesses with respect to feature selection capability. Korthauer et al. [4] performed a benchmark on methods for controlling false discovery rate in repeated hypothesis testing, from which we evaluate the Benjamini-Hochberg and q-value methods. However, these articles did not contain any numerical analyses in the context of right-censored data. More pertinent to our work is [5], where the authors compared fourteen filter methods for right-censored data. Considering mainly the prediction capability, the authors identified two competitive filtering methods, the variance filter and the correlation-adjusted regression survival scores (CARS scores) filter [6]. We have discussed CARS in the benchmark design section, but not the variance filter method as it is conceptually incompatible with our simulation studies.

In the context of right-censored data, Spooner et al. [7] evaluated prediction accuracy across a range of approaches, including regularized methods such as LASSO, ridge, and elastic net; CoxBoost alongside other machine learning algorithms; and two tree-based methods, including the random survival forest. The authors found CoxBoost to be the best in terms of prediction accuracy based on the concordance index.

In the benchmark design section, we discuss methods that are often used in analyzing high-dimensional data or that have been recommended by previous benchmarking articles, and have readily available software for implementation. The methods we consider are the least absolute shrinkage and selection operator (LASSO) [8, 9], adaptive LASSO (ALASSO) [10], elastic net (ENET) [11], CoxBoost (CB) [12], two random survival forest (RSF) [13] approaches, the Benjamini-Hochberg procedure (BH) [14], the q-value procedure (QV) [15], and the CARS [6] filter method recommended by [5]. We apply these methods to analyze synthetic datasets and compare their performance in terms of two feature selection metrics, the false discovery rate (FDR) and F1-score which combines FDR and the false negative rate; and three predictive metrics, the concordance index, Brier score, and root mean squared error of prediction defined in the benchmark design section. To our knowledge, this newly defined RMSE and other metrics have not been considered together in any of the benchmarking articles, nor has the diversity of our innovative synthetic data been achieved before in the context of right-censored data regression.

The results section details the outcomes of the study with accompanying figures and tables for all separate analyses and covers the most significant findings.

The discussion section provides brief coverage of additional methods that have been excluded from the study, additional details on the execution of the studies, and further commentary on the performances of the examined methods.

The unique contribution of this research are comparing several competitive methods in a wide variety of simulation settings, assessing their performance in terms of feature selection and prediction, and offering guidance to practitioners relying on the simulation results.

## Benchmark design

### Description of the analyzed methods

The analyzed methods can be broadly divided into embedded (model fitting) and filter methods. In an embedded feature selection method, feature selection is performed as a part of the model training process so that no extra step is needed for feature selection [2]. For embedded methods, we examine LASSO, ALASSO, ENET, CB, and RSF methods.

On the other hand, a filter method performs feature selection independently of any predictive model fitting. It uses statistical or mathematical criteria to rank and choose features before training a model. For filter methods, we examine the BH, QV, and CARS filter procedures.

### LASSO

The least absolute shrinkage and selection operator (LASSO) [8] is a well-known regularization method for estimating parameters in a high dimensional regression model. The method imposes a L1 penalty on the parameter, resulting in the “shrinking” of small parameter estimates to zero, and some shrinkage to non-zero estimates. When applying this L1 regularization to the partial likelihood, the regression parameter estimate is

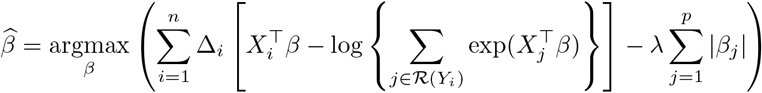

where ℛ (*y*) = {*r* : *Y*_*r*_ ≥ *y*} denotes the set of observations at risk of failure or experiencing failure at time *y* (the risk set), and *λ* denotes the L1 (LASSO) penalization term. In the R package `glmnet` [9, 16] that we use to implement the method, the optimal *λ* is estimated by 10-fold cross-validation to maximize the concordance of observations. When performing the LASSO, it is advised that all covariates are approximately on the same scale with similar means and variances.

### ALASSO

The adaptive LASSO (ALASSO) [10] is more general than the LASSO penalization method, where regularization is allowed to vary over the features. This adaptive regularization helps in selecting important features in the presence of correlation. The ALASSO log-hazard ratio vector estimate is

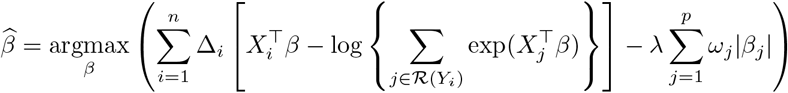

where *ω* = (*ω*_1_, *ω*_2_,···, *ω*_*p*_) is a vector of nonnegative *p*-dimensional known weights.

The goal of the inclusion of these weights is to increase the shrinkage of the parameter estimates corresponding to noninformative features and minimize the shrinkage of parameter estimates for significant features. Among several choices we set the weight 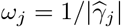, where 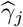 is the *j*-th estimated log-hazards ratio when the Cox proportional hazards model is fit to the data including all features with the ridge penalty (L2). A smaller value of 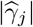 leads to higher penalty (or more shrinkage) for the *j*-th regression parameter. Once again, an optimal value for *λ* is obtained via minimizing a ten-fold cross-validation. We use the R package `Coxnet` [17] to obtain the ALASSO estimator.

### Elastic net (ENET)

ENET [11] is a generalization of the LASSO and Ridge regularized methods, where a mixture of L1 and L2 penalties are applied to the regression parameters. This encourages both complete shrinkage of noninformative features (L1 penalization) and more stable incomplete shrinkage of correlated features (L2 penalization). The ENET estimate is

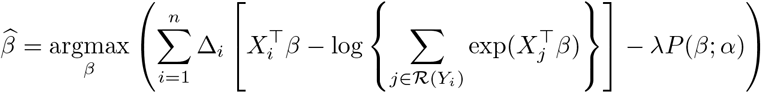

where

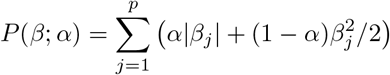

The parameter *α* ∈ [0, 1] determines the mixture of L1 and L2 penalties, and may be either estimated by cross-validation or fixed *a priori*. We fix *α* to 0.5. However, we estimate *λ* through ten-fold cross-validation. The R package `glmnet` [9, 16] is used to perform the ENET.

### CoxBoost (CB)

CB [12] is a method of estimating the regression parameters of the proportional hazards model (1) using a regularization and gradient boosting method. Although a machine learning technique is used in the estimation, CB is not a model-free approach. Suppose that *β*^(*t*)^ and 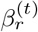 are the parameter vector and its *r*^*th*^ component at the *t*^*th*^ iteration. Define *η*_*r*_ = (*H*_*r*_(*β*^(*t*)^) + *λ*)^−1^*G*_*r*_(*β*^(*t*)^), where

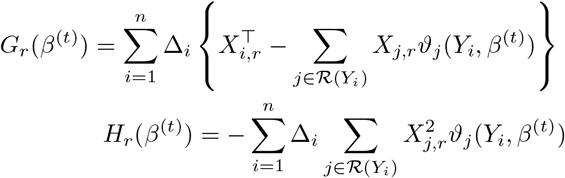

represent the gradient and double derivatives of the partial log-likelihood function with respect to the *r*th component of *β*-vector with

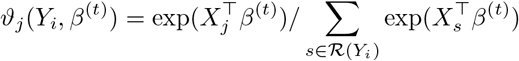

and *λ* is the penalty term to avoid overfitting. Define

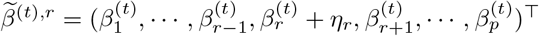. Determine

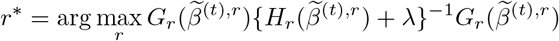

This index *r*^∗^ maximizes the squared scaled gradient, which is somewhat equivalent to maximizing the log-partial likelihood [18]. Then set 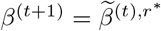, for *t* = 1, ···, *M*, where *M* denotes the number of boosting steps. In the accompanying R package `CoxBoost` [19], the default choice of *λ* is nine times the number of failures in the data, and the default for *M* is 100. We use these default hyperparameter values for all studies.

### Random survival forest (RSF)

The random forest, proposed by Breiman [20], seeks to capture the strengths of the nonparametric decision tree while reducing the overfitting that commonly occurs when fitting a single tree model. To accomplish this, a technique known as bootstrap aggregation (“bagging”) is employed, where many bootstrapped samples of the original data are used to fit weak learner decision trees which are then aggregated (normally through averaging) to form a single strong learner. Ishwaran et al. [13] extended this idea to censored survival data and provided a corresponding R package [21]. The main steps to fit a RSF are as follows.

1. Draw *B* (default 500) bootstrap samples from the original data, each excluding approximately 37% of the data, called “out-of-bag” (OOB) data.
2. Grow a survival tree for each bootstrap sample. For each node, randomly select 403 candidate variables. The node is split using the candidate variable and location that maximizes mean survival difference between child nodes.
3. Grow the tree to full size under the constraint that every terminal node should have at least eight failures.
4. Calculate the survival function for each terminal node (which is briefly described below).

At every terminal node *h* of all trees *b* in the forest, a Kaplan-Meier survival function is approximated using all (even OOB) observations that lie within *h*. Let 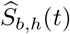 denote the Kaplan-Meier estimator at node *h* of tree *b*. To approximate the survival function for a new observation with feature vector *X*_0_, it is passed down all *B* trees until a terminal node *h*(*X*_0_) is reached. The arithmetic mean 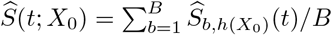 is the aggregated survival function estimate for this observation. The median of this estimate 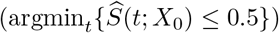 is used when predicting the survival time of this observation.

For a feature, minimum depth is the distance (or the number of steps) from the root node when the feature first appears in any given tree. Note that an important feature generally branches out the data closer to the root node than at further distanced nodes, so that important features are expected to be split upon early in a majority of trees in the forest. Once the RSF is fit, each feature’s average minimum depth is calculated over all trees. Features whose average depth is less than the mean of all average depths are selected as important features [22, 23].

Tuning parameters for the number of candidate variables at each node split and the minimum node size were chosen by a short simulation study, where optimal hyperparameters were estimated by minimizing out-of-sample error on 200 synthetic datasets. The ceiling of mean optimal hyperparameters (403 candidate variables, minimum node size of 8) are used in all studies. R code for this procedure is in `TuneRSF.R` of our repository. This *a priori* hyperparameter tuning was done to preserve computational resources in our large-scale simulation studies where RSF is already intensive, and may have resulted in some performance loss. We have performed an additional study that quantifies this loss in S1 Appendix, titled “Impact of RSF tuning parameters on performance.”

We chose to evaluate out-of-sample predictive metrics using observations that were unseen by the RSF during the training process (held-out data). Alternatively, out-of-sample predictive ability can be quantified by using OOB data, which acts as a proxy for out-of-sample data. For each observation, only the trees that did not include it during training (its OOB trees) are used to generate predictions. We have performed a simulation study comparing these procedures in S1 Appendix, titled “Comparing held-out and out-of-bag predictive metrics in RSF.”

### Screened random survival forest (sRSF)

We first ran univariate Cox regression of the response on each individual covariate. Any covariate with a p-value greater than 0.1 was excluded. The remaining covariates were then used as inputs to the RSF model, which was fitted as described previously. For evaluating feature selection capability of the method, features were selected using the minimum depth criteria after fitting the RSF.

### Benjamini-Hochberg procedure (BH)

The Benjamini-Hochberg (BH) [14] and q-value (QV) [24] procedures are two variable screening methods that have been proposed to control the rate of false discovery during repeated hypothesis testing on high-dimensional data. Both of these procedures are applied on *p* p-values arising from *p* hypothesis tests, one for each predictor through univariate Cox regression, and the methods return FDR-adjusted measures, which are used for feature selection. Without these adjustments, using standard *p*-values and independent *α*-level significance tests results in approximately *α* · (*p* − *s*) type-1 errors (false discoveries), where *s* denotes the number of true significant features. The details for these methods are outlined below.

Suppose that *p* null hypotheses *H*_1_, ···, *H*_*p*_ correspond to p-values *P*_1_, ···, *P*_*p*_ such that *P*_1_ ≤ *P*_2_ ≤ ··· ≤ *P*_*p*_. Let

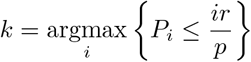

for a given desired FDR *r*. BH advises rejecting *H*_1_, *H*_2_, ···, *H*_*k*_. In our context, *H*_*i*_ : *β*_*i*_ = 0, where *β*_*i*_ is the log-hazards parameter corresponding to the predictor with associated *p*-value *P*_*i*_ for the Cox proportional hazards model fit to the *i*-th predictor. Rejecting *H*_*i*_ is analogous to considering the *i*-th predictor to be significant. This procedure ensures that the expected FDR is smaller or equal to *r* when the *p*-values are independent [14] or positively dependent [25]. For arbitrary dependence (e.g., features might be correlated in an arbitrary fashion), a more conservative threshold has been proposed [25], which is not considered here.

We then used the selected features in a multivariate Cox regression model and calculated predictive metrics based on the fitted model. The R package `mutoss` [26] was used to perform BH.

### Q-value (QV)

In this procedure, for each test a q-value is calculated, which is the minimum FDR at which that test would be called significant [24]. For that calculation, the proportion of true null hypotheses *π*_0_ is estimated from the data by the “smoother” method provided in the `WGCNA` [27] package implementation of QV. Assume *p* hypotheses are being tested. Corresponding to p-value *P*_*i*_, the q-value is

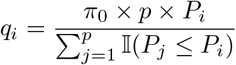

where 𝕀 (•) is an indicator function taking on one or zero if the statement • holds or not. If we decide to reject *H*_*i*_, the expected FDR among all features with q-values less than or equal to *q*_*i*_ is *q*_*i*_. The q-value method assigns an FDR estimate to each individual test (analogous to a p-value), allowing identification of significant features across a range of FDR thresholds. In contrast, BH requires repeating the significance decision process for each chosen FDR cutoff. QV is generally more powerful than BH.

For prediction, the selected features are then used to fit in a multivariate Cox regression model to the time-to-event data.

### Correlation-adjusted regression survival (CARS) scores

The CARS filter is an extension of the correlation-adjusted regression scores filter from Zuber et al. [28] adapted to right-censored data. Welchowski et al. [6] defined the vector of CARS scores as

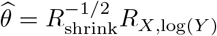

where *R*_shrink_ ∈ ℛ^*p*×*p*^ is a shrinkage estimator of the correlation matrix of *X*, and *R*_*X*,log(*Y*)_ ∈ ℛ^*p*×1^ estimates the correlations of the columns of *X* with log(*Y*) with inverse-probability-of-censoring weighting to counteract biasing by censored observations. The general inverse-probability-of-censoring weight of the *i*-th observation is defined as

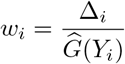

where 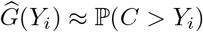 approximates the censoring process, considering *C* as a random variable independent of *T*_*i*_ and *X*_*i*_, i.e. the censoring process is noninformative. The matrix *R*_shrink_ is used in place of the correlation matrix between the columns of *X* to simplify computation of the inverse square root for large *p*. 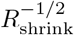 is found using singular value decomposition of *R*_shrink_. For further details, see Welchowski et al. [6]. Predictors with CARS scores further away from zero are considered more strongly associated with time-to-event than those with the scores closer to zero, and their signage determines the direction of the association. The corresponding R package [29] advises selecting features with the top 5% of |*θ*| as important predictors. Since the 5% could be very small or quite large, depending on the number of features, we employed two techniques on |*θ*| to identify important features.

First, order the absolute value of |*θ*|, and call the ordered values 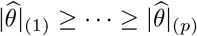. Let *L* be the fitted straight line through the points 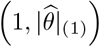 and 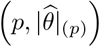, and define *D*(*i, L*) as the shortest (perpendicular) distance between the line and the point 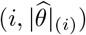. The elbow point is

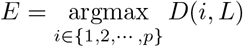

Features having 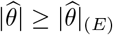 are considered to be important features. We refer to this process of elbow point estimation as maximal Euclidean distance (MED). The MED procedure tends to select more features. Therefore, we also propose an ad-hoc approach to determine this elbow point. For all *k* in {2, 3,…, *p* −1}, define the sets 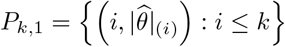 and 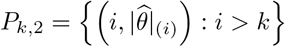. Fit two least-square linear regressions, one to points in *P*_*k*,1_ and one to *P*_*k*,2_, and collect the residuals. Let *e*_1:*k,i*_, *i* = 1, ···, *k* be the residuals from the first regression and *e*_(*k*+1):*p,i*_, *i* = *k* + 1, ···, *p* from the second regression. We define the elbow point as

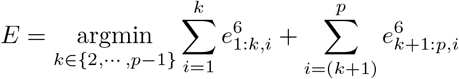

Using lower powers than 6 gives a larger estimate of *E*, which we opt against. We refer to this procedure for estimating the elbow as the minimal sextic residuals (MSR).

Fig 1 shows an example of the elbow point estimates from these two methods on absolute CARS scores calculated on a real bladder cancer dataset.

**Fig 1.**
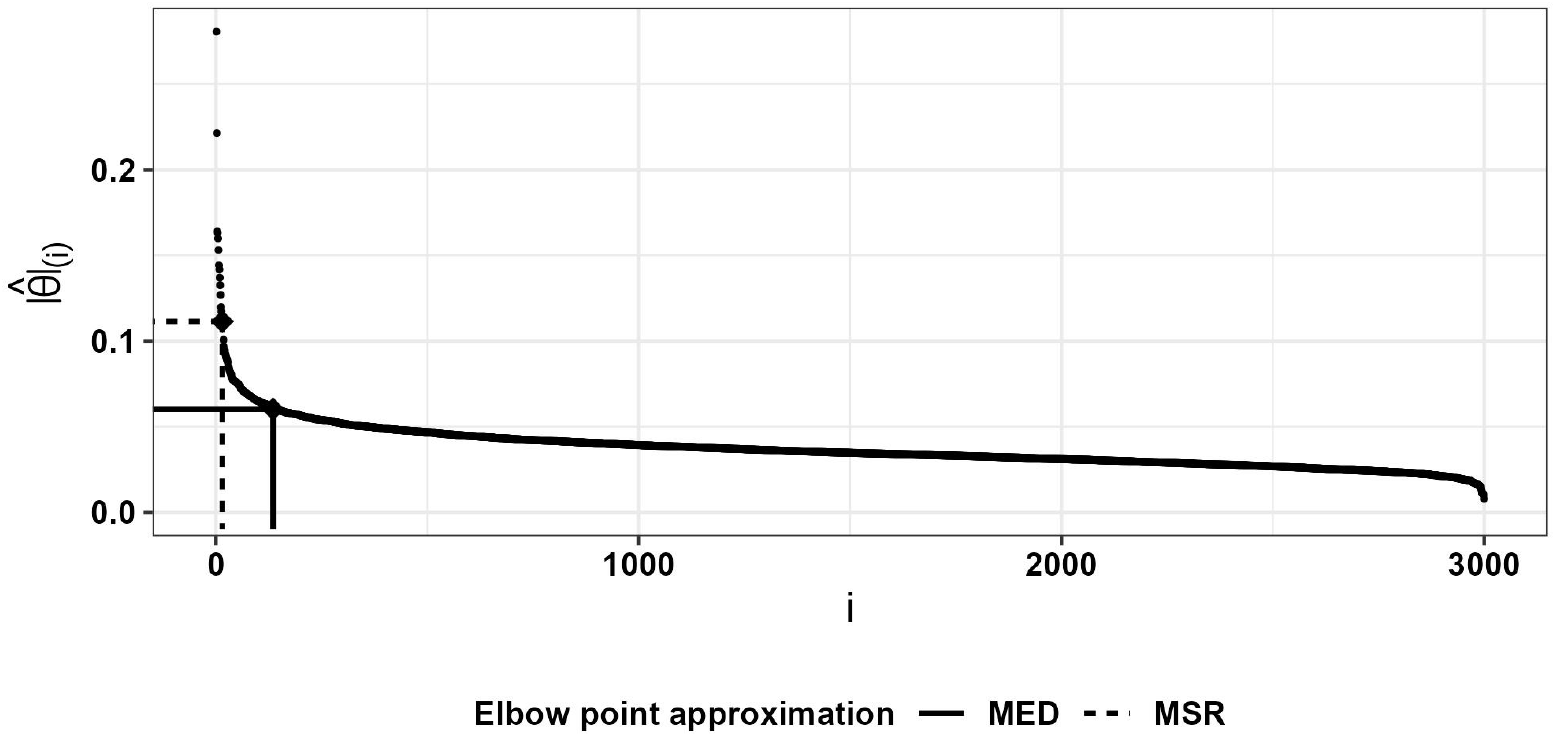
Elbow point estimation using the MED and MSR methods on the plot of decreasing absolute CARS scores for 3,000 features from the bladder cancer cohort (TCGA-BLCA).

We considered both methods for elbow point estimation in selecting predictors after computing CARS scores. The selected features are used to fit a multivariate proportional hazards model.

A brief summary table of the examined methods’ characteristics is provided in Table 1. Recall that embedded methods combine the steps of feature selection and model-building while filter methods perform these steps sequentially, and that parametric methods assume that the underlying distribution of survival times follows (1), so that the regression parameter vector *β* is estimated by the method.

**Table 1.**
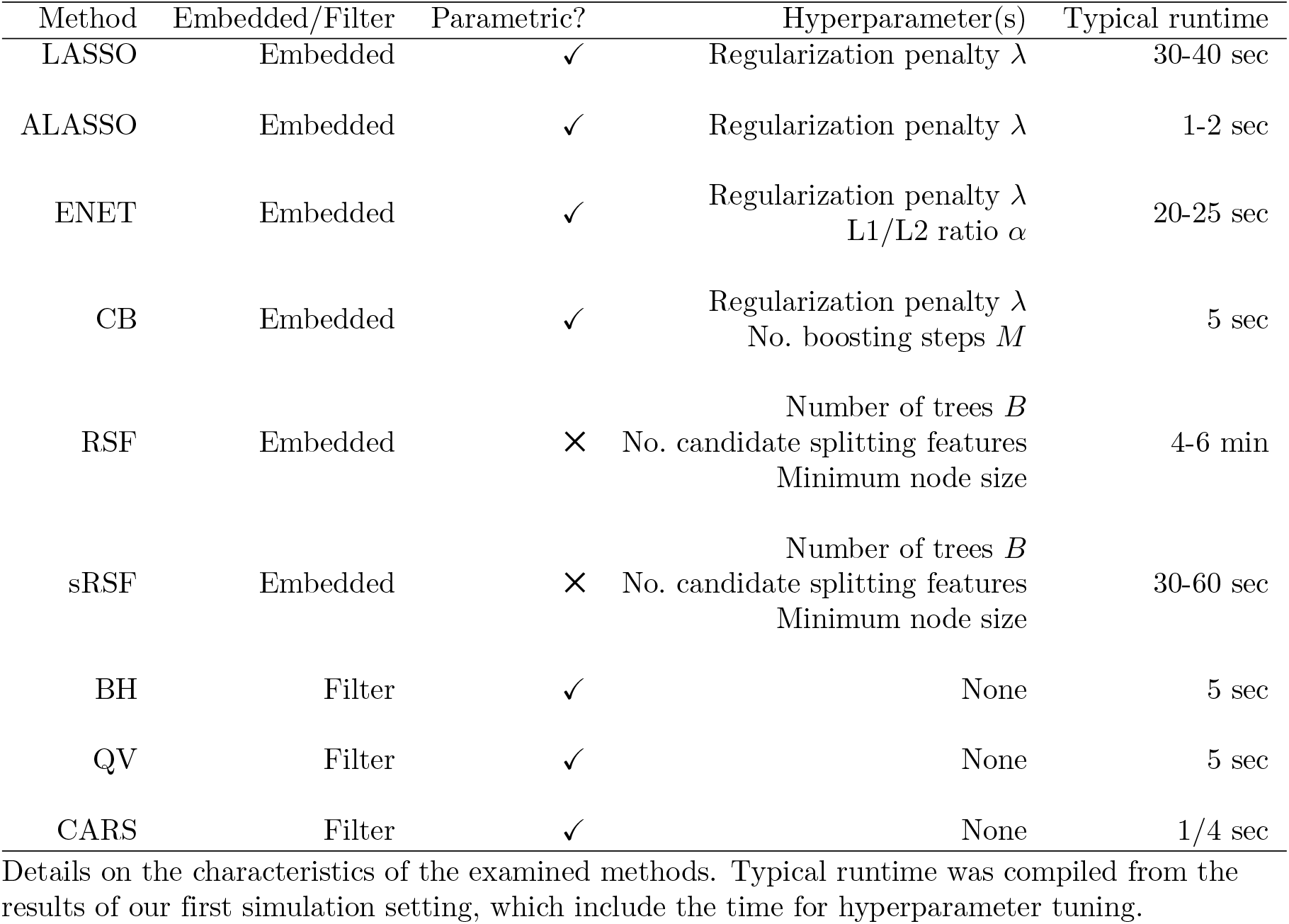
Methods at a glance.

| Method | Embedded/Filter | Parametric? | Hyperparameter(s) | Typical runtime |
| --- | --- | --- | --- | --- |
| LASSO | Embedded | ✓ | Regularization penalty $\lambda$ | 30-40 sec |
| ALASSO | Embedded | ✓ | Regularization penalty $\lambda$ | 1-2 sec |
| ENET | Embedded | ✓ | Regularization penalty $\lambda$<br>L1/L2 ratio $\alpha$ | 20-25 sec |
| CB | Embedded | ✓ | Regularization penalty $\lambda$<br>No. boosting steps $M$ | 5 sec |
| RSF | Embedded | ✗ | Number of trees $B$<br>No. candidate splitting features<br>Minimum node size | 4-6 min |
| sRSF | Embedded | ✗ | Number of trees $B$<br>No. candidate splitting features<br>Minimum node size | 30-60 sec |
| BH | Filter | ✓ | None | 5 sec |
| QV | Filter | ✓ | None | 5 sec |
| CARS | Filter | ✓ | None | 1/4 sec |

Details on the characteristics of the examined methods. Typical runtime was compiled from the results of our first simulation setting, which include the time for hyperparameter tuning.

### Simulation study

#### Synthetic data structure (setting-I)

The *p*-dimensional feature vector *X* was simulated from Normal(0, Σ), where Σ = (1 − *α*)*I*_*p*_ + *α*1_*p*_1^⊤^*p*, where *I*_*p*_ and 1_*p*_ respectively denote the identity matrix of order *p* and the *p*-vector with all entries being one. We considered two different values of *α*, 0 (the independent features case) and 0.5 (the correlated features case). For every *X*, we simulated the time-to-event *T* from the exponential distribution with scale exp(*X*^⊤^*β*). Then censoring time *C* was simulated from Uniform(*Q*(0.5), *Q*(0.9)), where *Q*(*r*) denotes *r*^*th*^ sample quantile of the *T* vector. Then observed time was set to *Y* = min(*T, C*) and censoring indicator Δ = 𝕀 (*T < C*). The procedure resulted in about 15% right-censored observations. The sample size for every simulated dataset was *n* = 300 and *p* was 1000.

We considered three different levels of sparsity (the percentage of nonzero components of the regression vector *β*), *s* = 2%, 5%, and 10% and set *β*_1_ = ··· = *β*_*ps/*2_ = *γ, β*_*ps/*2+1_ = ··· = *β*_*ps*_ = −*γ, β*_*ps*+1_ = ··· = *β*_*p*_ = 0, for *γ >* 0. We took three different values for *γ*, 0.5 for weaker signals, 1, and 2 for moderate to strong signals. The simulation design was somewhat similar to that of Zhu et al. [30].

For every unique combination of *α, s, γ*, we generated 200 datasets through the process outlined above. In every dataset, 200 of the 300 total observations were analyzed by each of the methods described earlier, and the remaining 100 were used to evaluate the out-of-sample predictive ability of each resulting model. A data-flow diagram for this study is presented in Fig 2.

**Fig 2.**
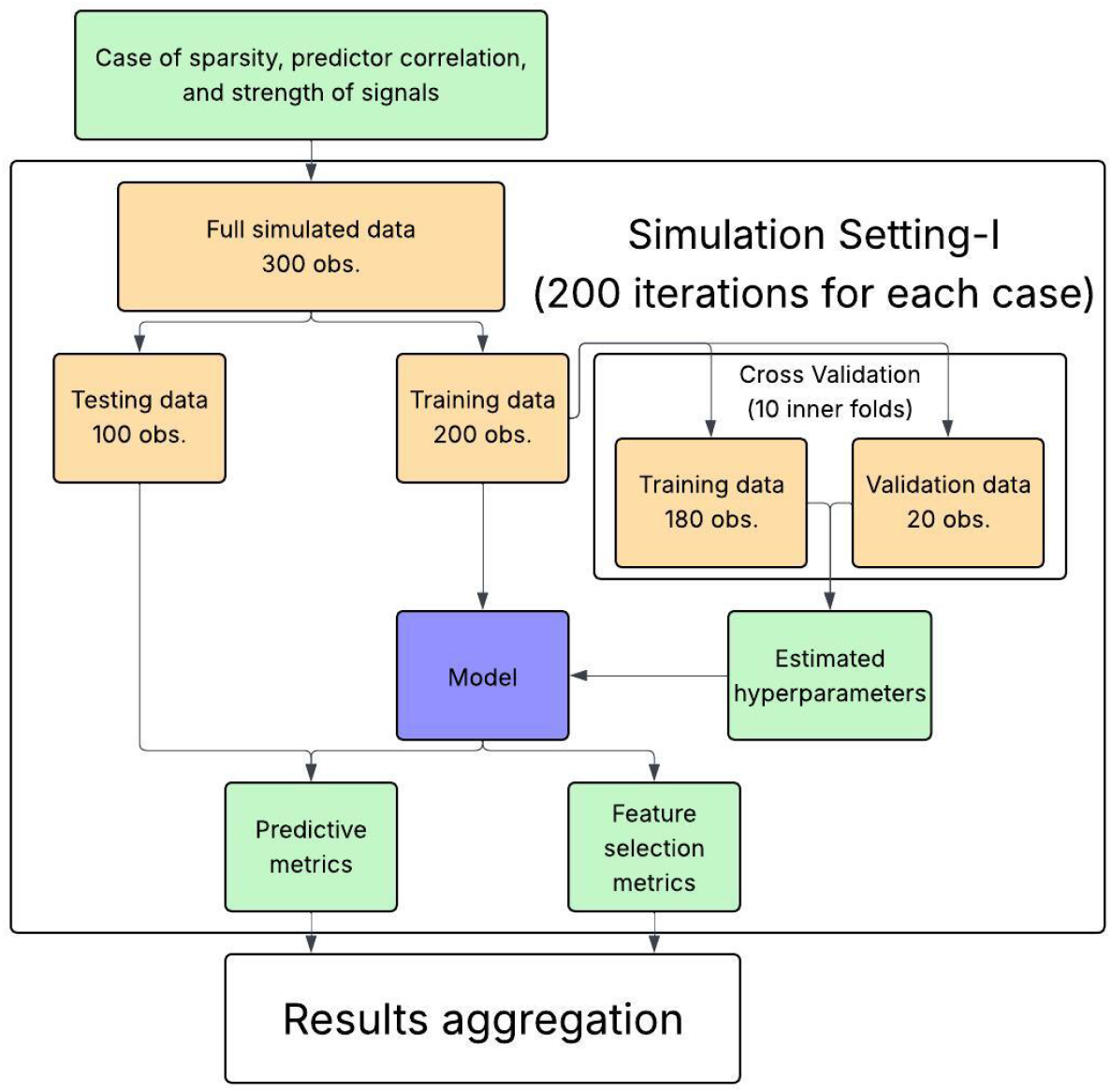
Data-flow diagram of simulation setting-I. 18 unique cases of data characteristics are considered, each using *s* ∈ {0.02, 0.05, 0.1} (sparsity), *α* ∈ {0, 0.5} (predictor correlation), and *γ* ∈ {0.5, 1, 2} (strength of true signals). Methods that do not require hyperparameter tuning do not undergo the cross-validation step.

#### Synthetic data simulation imitating the bladder cancer data (setting-II)

In this setting, we simulated data mimicking a real bladder cancer cohort from The Cancer Genome Atlas (TCGA-BLCA). The sample size was set to *n* = 423 (the real data sample size) and the number of features was 3,000, which was a pre-screened subset of a larger number of mRNAs standardized to mean 0 variance 1. We set the true regression parameter *β* to 10 times the log-hazard estimates of the real data using the CoxBoost method, which had 53 nonzero coefficients out of the 3,000 coefficients. All nonzero coefficients were between 0.15 and 0.9 in absolute value. The time-to-event was simulated from the Weibull distribution with shape 2 and scale 300 exp(−*X*^⊤^*β/*2), i.e., *P* (*T > t*|*X*) = exp{− (*t/*[300 exp(−*X*^⊤^*β/*2])^2^}, so that the median survival time was approximately 250. Censoring time *C* was randomly simulated from Uniform(*Q*(0.5), *Q*(0.9)), where *Q*(*r*) is the *r*-th sample quantile of *T*. Each method was applied for analyzing the data (*X*_*i*_, *Y*_*i*_, Δ_*i*_), *i* = 1, ···, 423, where *Y*_*i*_ = min(*T*_*i*_, *C*_*i*_) and Δ_*i*_ = 𝕀 (*T*_*i*_ *< C*_*i*_). We generated 200 datasets with the above process for this setting of simulations. In each dataset, 43 (approx. 10%) observations were chosen at random to be for evaluating out-of-sample predictive performance. The remaining observations were used for training of all methods. A data-flow diagram for this setting of simulations is shown in Fig 3.

**Fig 3.**
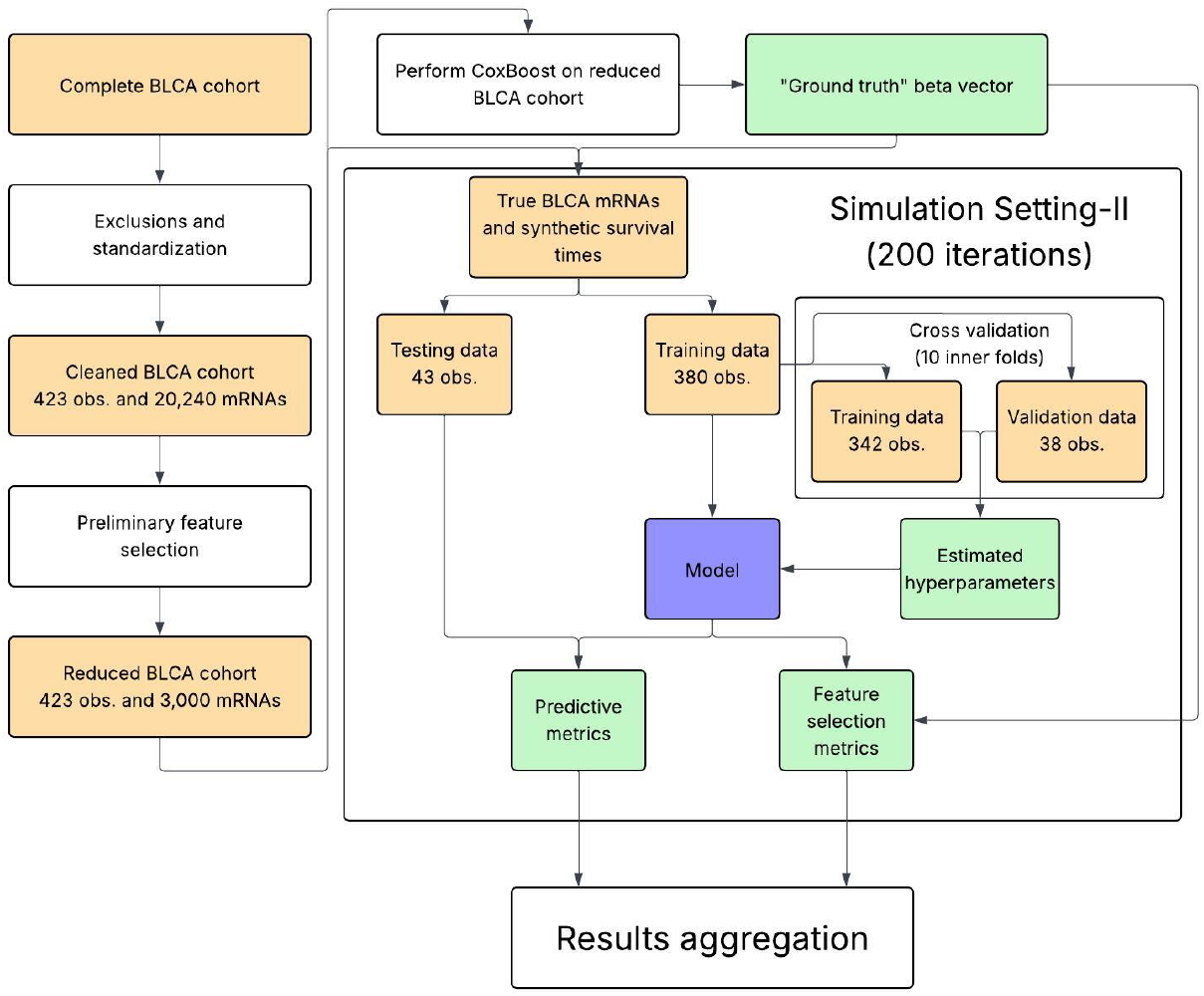
Data-flow diagram of simulation setting-II. The TCGA-BLCA cohort cleaning and the preliminary feature selection step are discussed in the real data analysis background. Methods that do not require hyperparameter tuning do not undergo the cross-validation step.

#### Performance evaluation metrics

##### Feature selection metrics

We employed FDR = FP*/*(TP + FP) and F1-score = TP*/*(TP + 0.5(FP + FN)) to evaluate feature selection performance, where TP, FP, and FN denote the number of true positives, false positives and false negatives, respectively. Both FDR and F1-score lie between zero and one. FDR is an increasingly important metric to consider in the field of bioinformatics [4], as falsely classifying irrelevant variables as significant can result in a major waste of resources in pursuit of false leads in biological research. We also seek to succinctly evaluate the precision = TP*/*(TP + FP) and recall = TP*/*(TP + FN) of the methods, so we employ F1-score, which is their harmonic mean. Optimal methods will have a low FDR and a high F1-score.

#### Predictive capability metrics

##### Concordance index (CI)

We evaluated the predictive ability of the methods with out-of-sample CI [31]. Two observations are considered comparable if one has experienced an event before the other either experiences an event or is censored. A comparable pair of observations is concordant if the observation that experiences an event first is predicted to have a shorter survival time (or equivalently be at greater risk), and otherwise the pair is discordant. CI is defined as the proportion of concordant pairs among the comparable pairs.

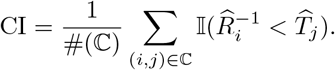

Here ℂ is the set of comparable observation pairs (*i, j*) such that *Y*_*i*_ *< Y*_*j*_ and Δ_*i*_ = 1, and 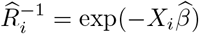 is the *i*-th observation’s (with covariate *X*_*i*_) inverse risk estimate where 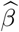 is the method’s log-hazard ratio vector estimate when performed on the training data. Here #(·) denotes set cardinality. For the nonparametric RSF and sRSF, 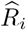 is the predicted survival time from the ensemble of decision trees. CI can take on values between 0 and 1 inclusive, where higher values show better predictive ability, and a score of 0.5 shows no improvement over randomly guessing each observation’s risk. We calculate CI using only out-of-sample observations.

##### Brier score

The Brier score [32] is analogous to the mean squared error between survival past a time point *t*_0_, 𝕀 (*T*_*i*_ *> t*_0_) and the predicted probability of survival past *t*_0_, 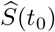. To account for event censoring, time-specific inverse-probability-of-censoring weighting of observations is used. The Brier score at time *t*_0_ is

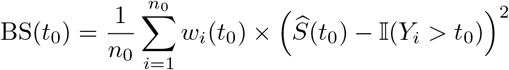

where 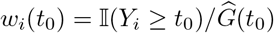 denotes the inverse-probability-of-censoring weight of the *i*-th observation at time *t*_0_, and *n*_*o*_ denotes the size of the out-of-sample data (test data). Again, 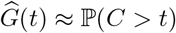 approximates the censoring process. In the first setting of simulations, we report the Brier score at *t*_0_ = 0.5, and *t*_0_ = 250 in the second setting, both of which are near the population medians of their respective studies. This metric is calculated on out-of-sample observations in all studies.

##### RMSE

We define the root mean squared error (RMSE) for the logarithm of the time-to-event as

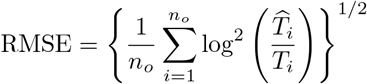

For parametric methods, 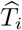 is the estimated median survival time under the Cox proportional hazards model fitted using only features with nonzero estimated regression coefficients. For RSF and sRSF, 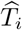 denotes the predicted survival time using the ensemble of trees. Here *n*_*o*_ denotes the size of the out-of-sample (testing) data. A lower value of RMSE indicates lesser difference between predicted and true event times, i.e. better predictive performance. This metric is calculated on out-of-sample observations. Notice that the true event times *T*_*i*_ are needed to compute the metric, making it exclusive to simulation studies.

In brief, CI quantifies how well observation risks are ranked, Brier score measures the gap between observed survival beyond a time point and the model’s predicted survival probability, and RMSE is the deviation between true event times and predicted event times.

### Real data analysis

#### Background

The Cancer Genome Atlas (TCGA) is a publicly available cancer genetics program containing genomic, epigenomic, transcriptomic, and proteomic data from cancer patients. The genomic data in particular have been used in benchmark studies (e.g., Bommert et al. [5]) and are generally accepted for demonstrating method efficacy on right-censored data. We used patients’ mRNA transcript abundance data from the bladder cancer (BLCA) cohort to illustrate how the examined methods can be performed on real data. We obtained the normalized (debiasing due to differences in sequencing depth or gene length) and log2 transformed (reducing skewness) mRNA data (HiSeqV2) prepared by the RSEM method [33] from the Xena Browser [34].

We obtained BLCA patients’ mRNA and time-to-event (days from diagnosis to patient failure) with censoring indicator and sex. We removed observations that experienced an event on the same day as diagnosis or contained missing values. Further, we removed all constant features. After these exclusions, we had 423 observations (patients), 308 of which were male and the remaining 115 female, and 20,240 mRNA features. Estimated Kaplan-Meier survival functions for male and female BLCA patients are shown in Fig 4, with corresponding log-rank test p-value for testing for differences between survival distributions. We also provide a histogram of the 20, 240(20, 240 − 1)*/*2 ≈ 204 million absolute pairwise correlations (APC) of the genes (Fig 5). To reduce to the high dimensionality and multicolinearity of the features, we define a preliminary feature selection (PFS) step.

**Fig 4.**
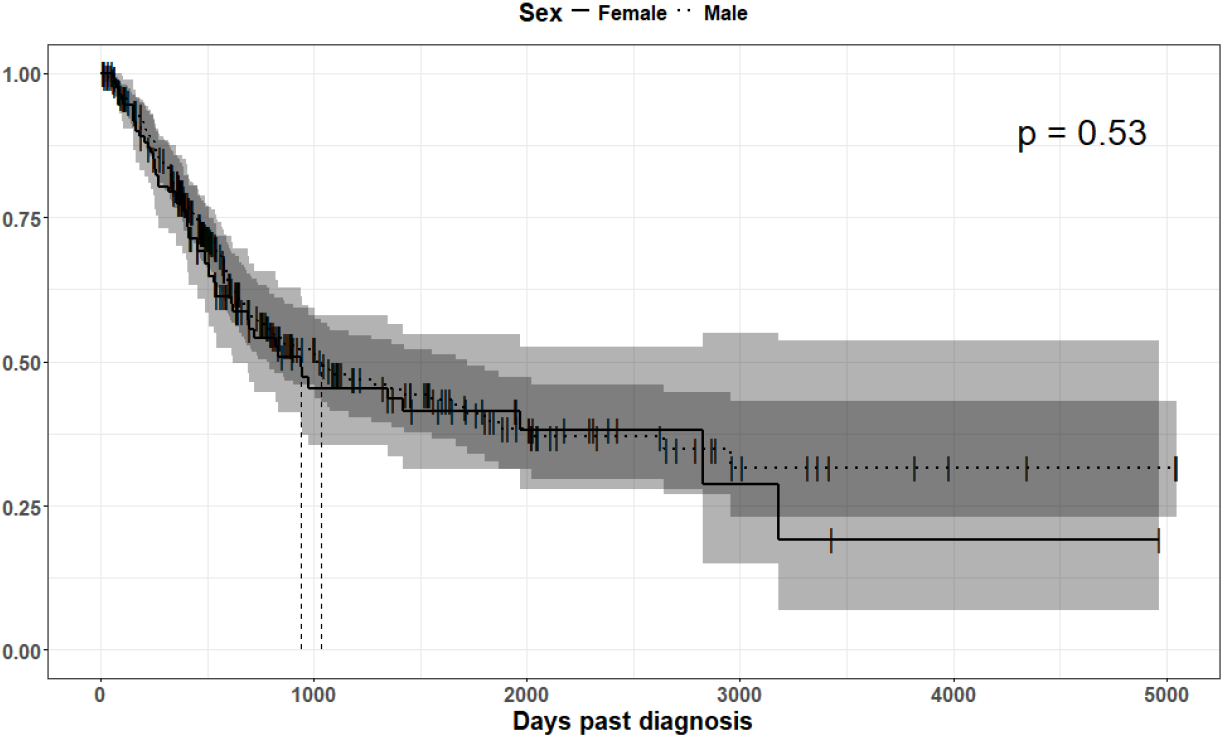
Estimated Kaplan-Meier estimator of the survival functions for BLCA cohort females and males. Shaded regions show 95% confidence intervals and vertical dashed lines are the estimated median survival times. The log-rank test’s p-value is inside the figure, which indicates no significant difference in survival distributions between the groups.

**Fig 5.**
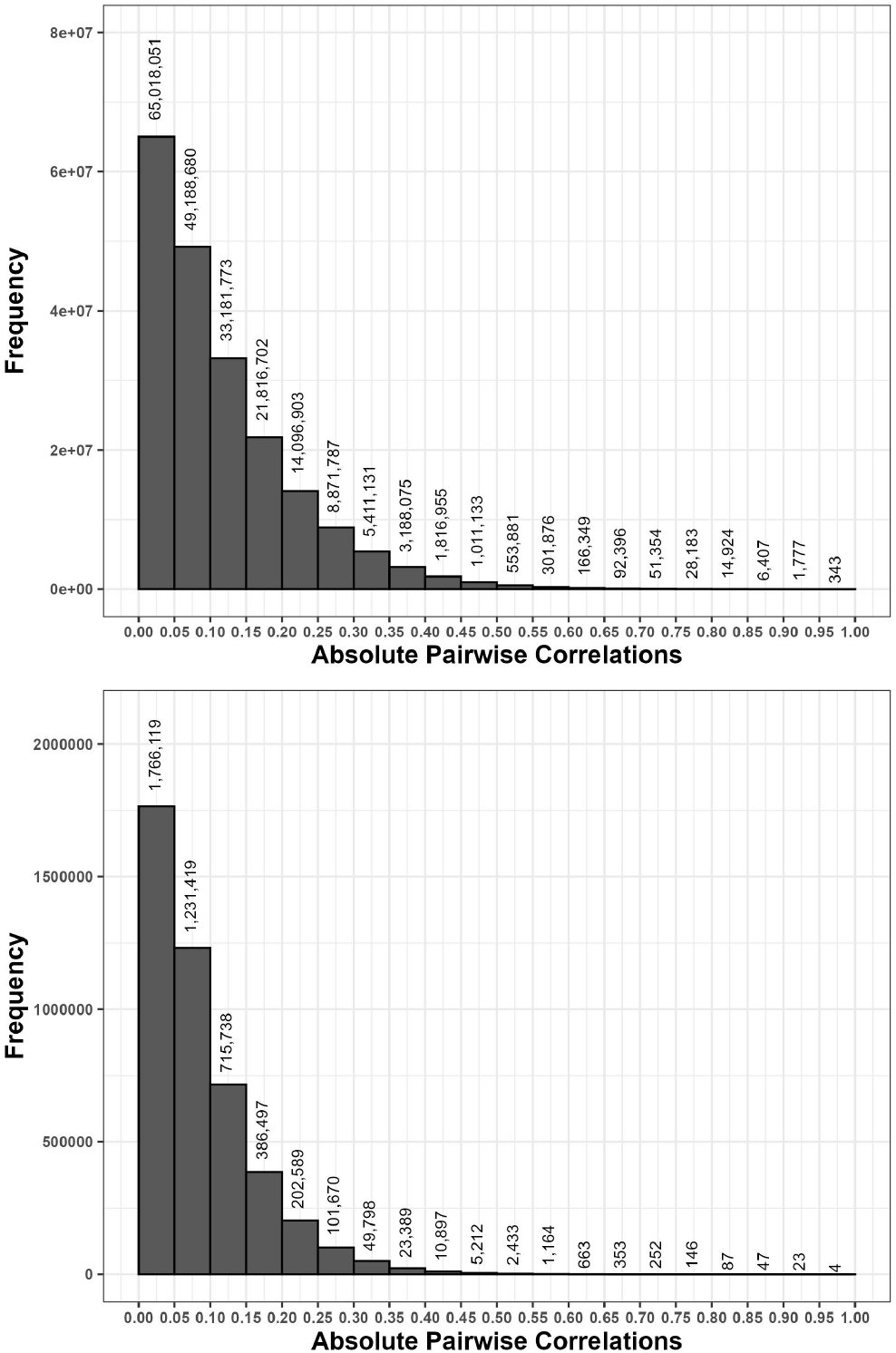
Absolute pairwise correlations (APC) before and after our preliminary feature selection (PFS). (Top) Histogram of APC between 20,240 mRNA features prior to PFS. (Bottom) Histogram of APC between the 3,000 features chosen by PFS. The number of observations within each bin are shown in the counts above each bar.

#### Preliminary feature selection (PFS)

The 20,240 mRNA features were standardized to have mean 0 variance 1 among all observations, and then CARS scores 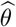 were calculated. The mRNAs corresponding to the 3,000 greatest values of 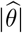 were kept and used by all methods in the real data analysis and setting-II of our simulations. CARS was chosen for this process as it enjoys good computation scaling with respect to *p* and selects features that have strong marginal correlation to the outcome and lesser correlation with other features [6]. We performed PFS once on all observations instead of per-fold so that methods select features each fold from a consistent feature set. Of the features kept by PFS, we examined existing literature to identify a feature subset whose elements likely contribute to differences in cancer progression and patient survival times. We found 30 features in our search, which are presented in Table 2 with corresponding literature citations.

**Table 2.** List of 30 mRNA features likely associated with differences in cancer patient survival time found within existing literature.

| Known driver | Existing literature | Known driver | Existing literature |
| --- | --- | --- | --- |
| ATG5 | Xu et al. [35] | DLEU1 | Li et al. [36] |
| FGF5 | Long et al. [37] | FGF14 | Su et al. [38] |
| FGF22 | Furuta et al. [39] | FGFRL1 | Tai et al. [40] |
| FOXA3 | Wang et al. [41] | FOXF1 | Bian et al. [42] |
| FOXF2 | Zheng et al. [43] | FOXG1 | Chen et al. [44] |
| FOXI1 | Onodera et al. [45] | FOXK2 | Kang et al. [46] |
| FOXL2 | Llana et al. [47] | FOXN4 | Zhang et al. [48] |
| FOXP3 | Szylberg et al. [49] | FOXR1 | Liao et al. [50] |
| IGF1R | Elvira et al. [51] | IGF2BP1 | Müller et al. [52] |
| MMP8 | Juurikka et al. [53] | MMP19 | Yu et al. [54] |
| MMP20 | Mok et al. [55] | MMP27 | Eichberger et al. [56] |
| PLA2G3 | Ray et al. [57] | PLA2G4A | Qian et al. [58] |
| PLA2G4D | Recuero et al. [59] | PSMD4 | Ma et al. [60] |
| PSMD13 | Li et al. [61] | SQLE | Zhao et al. [62] |
| TOP1 | Pfister et al. [63] | TOP3B | Al Mahmud et al. [64] |

A literature search was performed of the 3,000 mRNA features remaining after performing PFS. We included articles that performed laboratory studies or statistical analyses to conclude that variation in the corresponding mRNA’s expression contributed to differences in survival time.

### Additional studies concerning PFS

Three additional studies were performed that examine the quality of PFS and its downstream effects in the real data analysis, all described in S1 Appendix.

To examine the downstream effects of PFS, we evaluated the embedded methods’ differences in predictive accuracy and runtime in the real data analysis when performed using either the size 20,240 feature set (no PFS) or the reduced set of 3,000 features (with PFS). This study is titled “Effect of PFS on embedded methods’ performances in the real data analysis.”

We also examined the overall quality of the PFS step through two studies. Firstly, we observed the relationship between the signal-to-noise ratios of truly significant features (for which |*β*| is a proxy) and their retention rates when PFS is performed through a simulation study titled “Retention of true signals by PFS under varying signal-to-noise ratios.” Secondly, we performed a separate study that examines PFS with the real survival times to determine how sensitive the procedure is and if/how the downstream modeling may have been affected by its application on the full data instead of per-fold, titled “Sensitivity analysis of PFS.”

### Analyses techniques

Let ℳ denote the set of 3,000 mRNA features kept by PFS, and ℳ_0_ ⊂ ℳ be the subset of 30 true signals identified within existing literature. We randomly partitioned the 423 observations into 10 approximately equally sized and mutually exclusive folds. Every method was performed 10 times in this analysis, each time using the observations from 9 of the 10 folds for training. All features were shifted and scaled to have mean 0 variance 1 among the observations in the training folds. This process of repeatedly performing a method on a subset of all observations and evaluating predictive performance on the remaining observations (the outer loop) and performing hyperparameter cross-validation within the training set (the inner loop) during each iteration of the outer loop is known as nested cross-validation. We use this process to achieve replications in lieu of the ability to generate new datasets. The sets of selected features from the *i*-th model (denoted *S*_*i*_ ⊂ ℳ for *i* ∈ {1,···, 10}) were recorded, and out-of-sample CI and Brier score at 365 and 1,000 days post-diagnosis were calculated using the fold of observations left out of the training. For quantifying feature selection capability, we recorded #(*S*_*i*_) and #(*S*_*i*_ ∩ ℳ_0_) (i.e. the total number of selected features and the number of selected features among the features identified within existing literature respectively) for *i* ∈ {1,···, 10}. These values are presented as “Selected mRNAs” and “True positives.” Since the uncensored event times of all observations are not known, we cannot compute RMSE. Further, we cannot compute FDR nor F1-score since knowledge of the exhaustive list of true signals is not present in this analysis of real data. We instead report feature selection stability with the Dice coefficient which is described later. A data-flow diagram for this individual study is shown in Fig 6.

**Fig 6.**
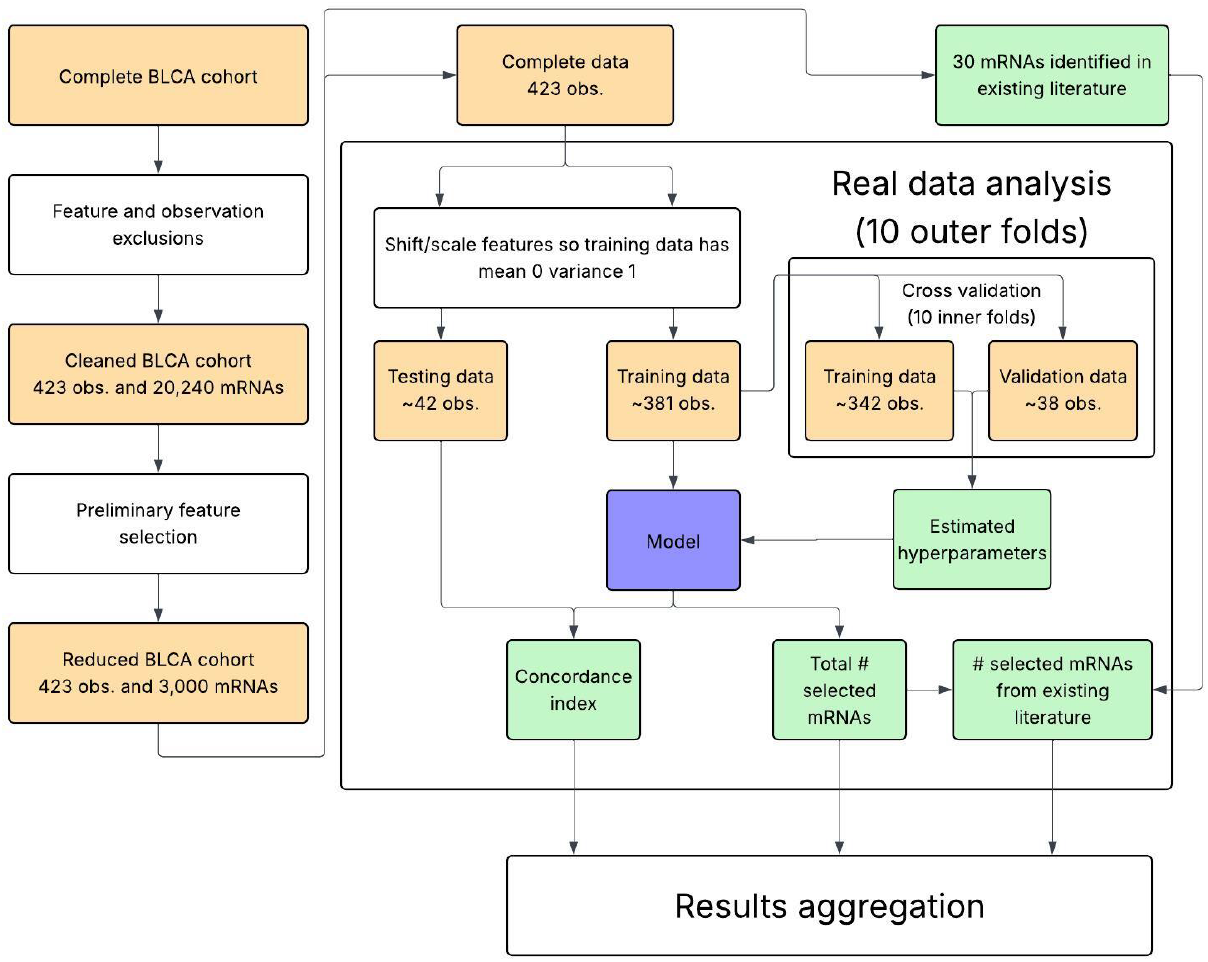
Data-flow diagram of the real data analysis. Nested 10-fold cross-validation of the full cohort was done so that methods were ran multiple times on the data and predictive performance was evaluated with out-of-sample observations. Methods that do not require hyperparameter tuning do not undergo the inner cross-validation step.

### Dice coefficient

The Dice coefficient [65] measures how “similar” two sets are by comparing the sizes of the individual sets with the size of their intersection. We calculate the Dice coefficient between all pairs of *S*_*i*_ and *S*_*j*_ where 1 ≤ *i < j* ≤ 10 for each method as

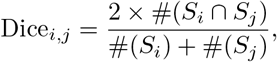

which varies between 0 when no features are in common between the sets and 1 when sets are identical. If one of the compared sets is the list of true signals then the resulting Dice coefficient is equivalent to F1-score. We report the median and inter-quartile range of the Dice coefficients for each method in the real data analysis results section.

## Results

### Simulation results (setting-I)

#### Feature selection performances

Boxplots of the F1-score and FDR for setting-I are shown in Fig 7. For each method, the boxplot displays the distribution of the metric across simulation replications, with the black horizontal bar indicating the median value. For example, when *α* = 0, *s* = 2%, and *γ* = 0.5, the median FDR for LASSO is about 75%.

**Fig 7.**
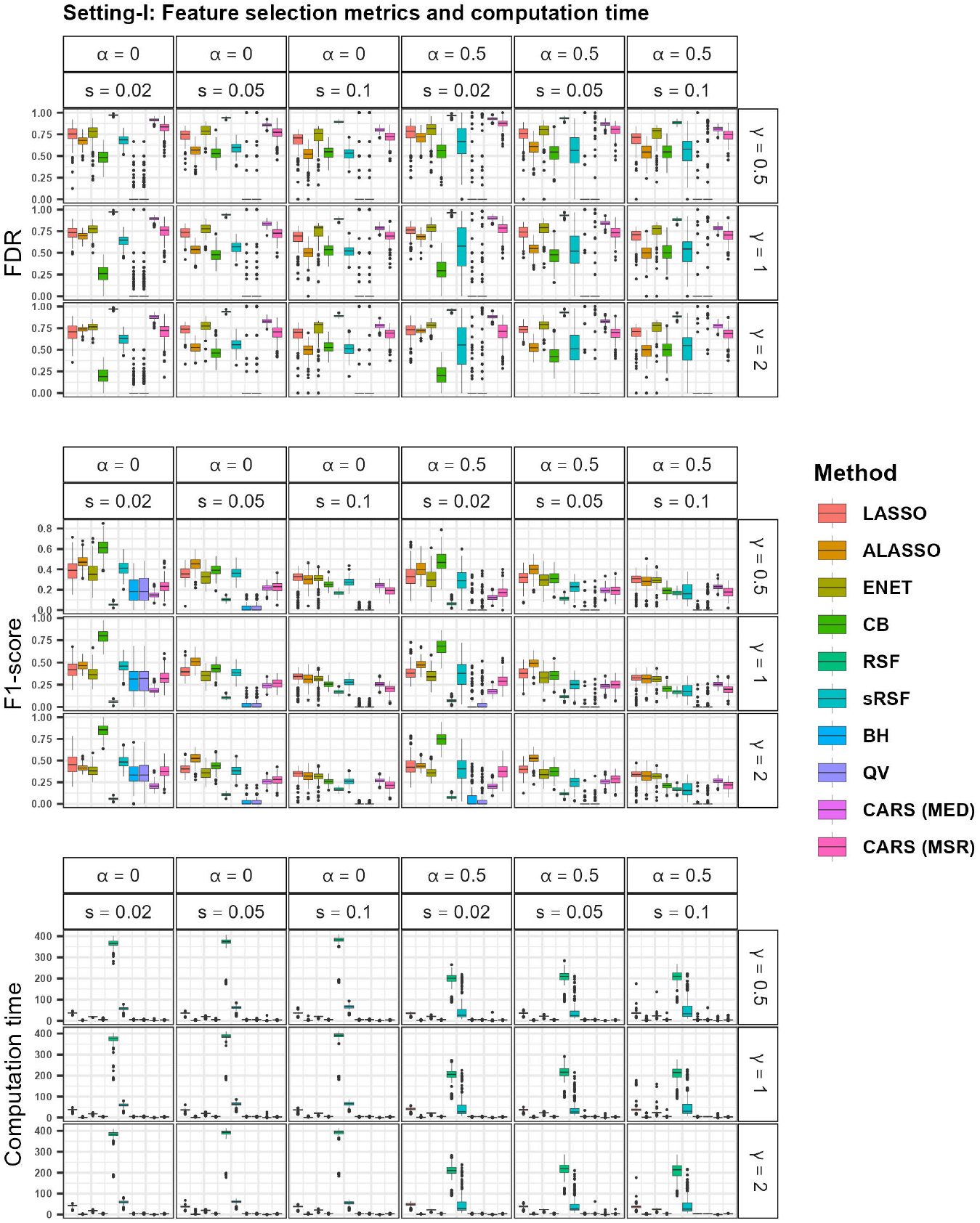
Boxplots of feature selection metrics and computation time in setting-I of the simulation studies. (Top) Boxplots of FDR. (Middle) Boxplots of F1-score. (Bottom) Boxplots of computation time, measured in seconds. Plots are faceted by data characteristics: horizontally by correlation between predictors and sparsity of informative features (*α, s*), and vertically by signal strength (*γ*).

Under most (*s, α, γ*) cases, the BH and QV procedures exhibit the lowest median FDR, while RSF shows the highest. For any given sparsity level, the FDR for ALASSO decreases notably as the sparsity *s* increases, especially when the signal strength *γ* is greatest. In contrast, increasing the correlation from 0 to 0.5 does not result in a noticeable change in FDR values. The FDR values from the sRSF procedure are substantially lower than those from RSF, indicating that the screening step helps reduce false discoveries. Unsurprisingly, MSR maintains lower FDR than MED when performing CARS due to its more conservative threshold for feature selection. Of the examined embedded methods, CB consistently has the lowest FDR when *s* = 2%, and is about tied with sRSF when *s* = 10%.

Note that a lower FDR indicates a method is more effective at avoiding the selection of unimportant features.

For sparsity 2% and 5%, ALASSO and CB show the highest F1-scores, with LASSO also performing competitively in the *γ* = 2 case. Overall, F1-scores increase with signal strength, while they decrease as the correlation among features increases. At the 10% sparsity level, LASSO shows the highest F1-scores, followed by ALASSO and CB. As expected, the BH and QV methods show the lowest F1-scores due to selecting very few features. Importantly, the F1-scores for sRSF are noticeably higher than those of RSF in most cases, indicating that the inclusion of the additional screening step improves the FDR and F1-score of the RSF method.

Note that a higher F1-score indicates a better method for detecting all relevant features while avoiding unimportant ones.

#### Predictive performances

Boxplots of CI, Brier score, and RMSE for setting-I are presented in Fig 8. A higher value of CI and lower values of Brier score and RMSE indicate better predictive capability.

**Fig 8.**
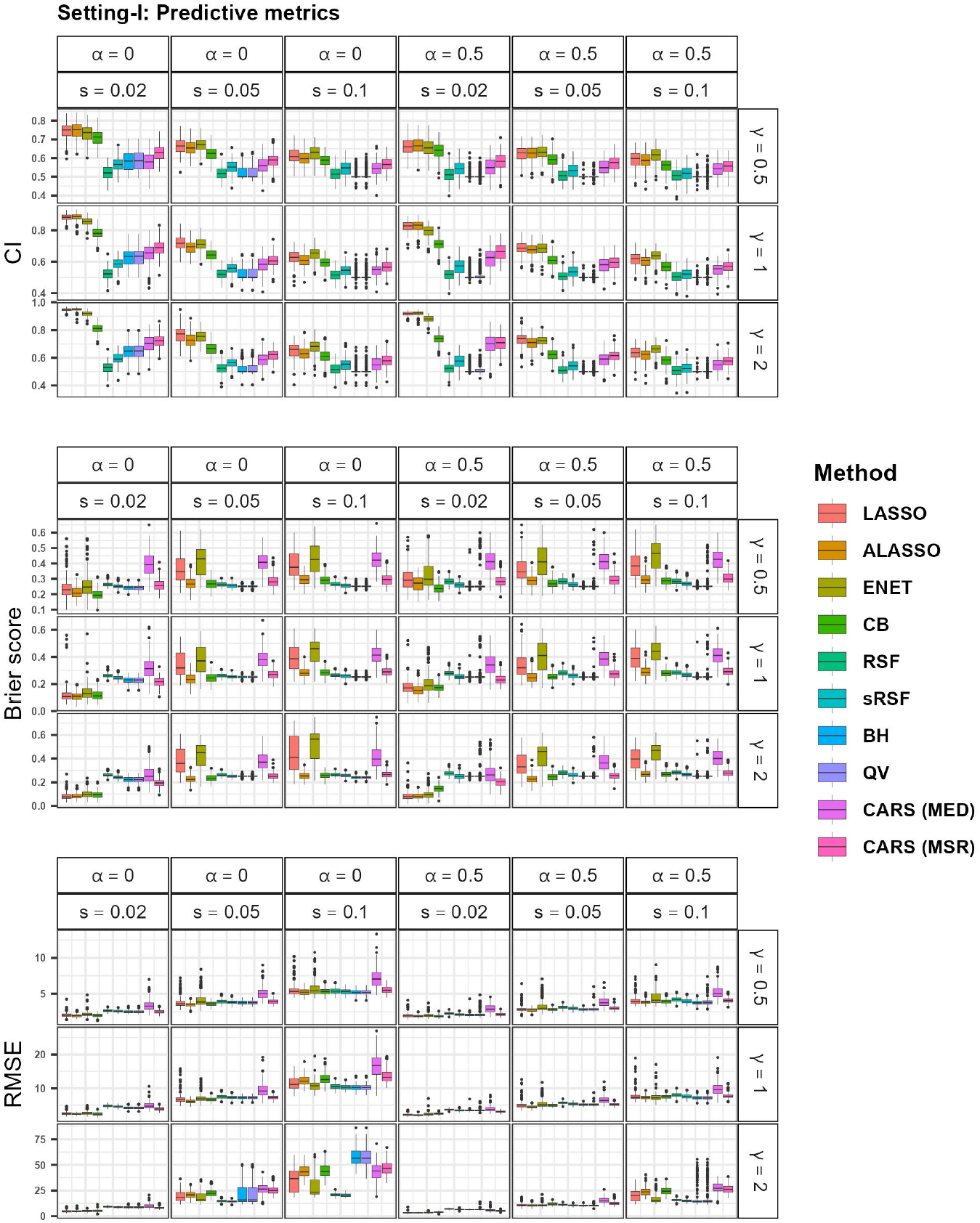
Boxplots of predictive metrics in setting-I of the simulation studies. (Left) Boxplots of CI. (Middle) Boxplots of Brier score. (Right) Boxplots of RMSE. Plots are faceted by data characteristics: horizontally by correlation between predictors and sparsity of informative features (*α, s*), and vertically by signal strength (*γ*).

In terms of CI, LASSO and ALASSO and ENET seem to perform the best across all scenarios. CB was generally around 4th in this metric. BH and QV maintain median CI of 0.5, suggesting these methods have no noticeable improvement over randomly guessing observation risk scores. Additionally, the MSR threshold outperforms MED when performing CARS. As expected, CI generally increases with *γ* and decreases with *s* and *α*.

Recall that methods with higher CI have superior predictions of relative risk between observations (i.e. observations with greater predicted risks generally experienced events prior to ones with lesser predicted risks).

When evaluating Brier score, the embedded methods generally outperformed the filter methods under the cases with low *s*. However, both classes of methods performed similarly when *s* = 10%. The ALASSO and CB were consistently the best performers among the embedded methods. CARS with the MSR threshold was the best of the filter methods for the case of low *s*, and BH and QV performed better when *s* was large.

Recall that Brier score quantifies the difference between predicted probability of an event occurring after a given time and whether an event was observed or not.

For smaller values of *s*, ALASSO is generally the best in terms of RMSE. In terms of CI and RMSE, sRSF performed better than RSF. In most cases, the MSR threshold produces lower RMSE than MED when performing CARS.

Note that methods with lesser RMSE more accurately estimate observation event times.

#### Computational time

Boxplots of methods’ computational time on our synthetic data (setting-I) are shown in Fig 7. Little variability in computation time was present as data characteristics changed, and relative rankings remained almost constant. RSF without screening was consistently the most computationally taxing, followed by the LASSO and sRSF. Interestingly, ALASSO is among the top performers. CARS also performed consistently well when using the MED threshold, but less so when using MSR.

#### Cumulative method rankings

To summarize each method’s overall relative performance in our first setting of simulations, we ranked the median metric values of all methods within all combinations of data characteristics, totaled the ranks over the combinations of characteristics, and finally ranked the totals. The resulting aggregate method rankings for each metric are presented in Table 3.

**Table 3.**
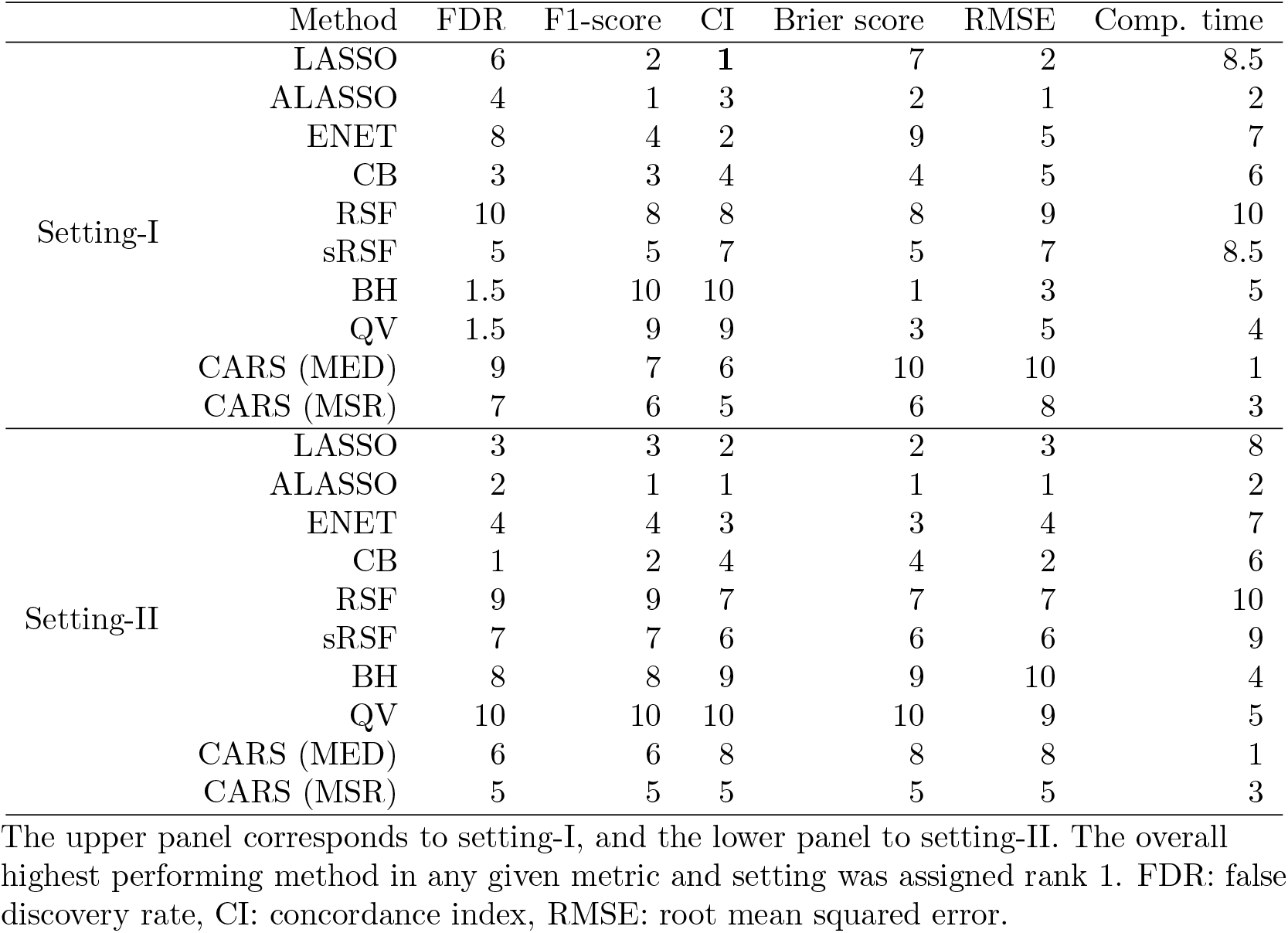
Aggregate relative method rankings in simulation settings I and II.

BH and QV consistently lead for FDR control, while ALASSO and LASSO achieved the highest F1-scores. For CI, LASSO and ENET generally outperformed, though BH had the best overall Brier score. LASSO and ALASSO also yielded the lowest RMSE. Regarding runtime, CARS with the MED threshold proved fastest.

A table containing the medians and inter-quartile ranges of metric values across each method’s 200 replications under each combination of data characteristics (*α, s, γ*) is provided in S2 Table. Each column of this table corresponds to a unique combination of data characteristics, and every row is an individual method and metric. The best median metric values for each case of data characteristics are boldfaced.

### Simulation results (setting-II)

#### Feature selection performances

A table of summary statistics (median and inter-quartile range) of all metrics for setting-II simulation studies are shown in Table 4. Well-performing methods have lower FDR, Brier score, RMSE, and computation time, and higher F1-score and CI.

**Table 4.** Summary statistics of setting-II of the simulation studies.

| Method | FDR | F1-score | CI | Brier score | RMSE | Computation time |
| --- | --- | --- | --- | --- | --- | --- |
| LASSO | 0.65(0.11) | 0.5(0.11) | **0.92(0.03)** | 0.08(0.04) | 1.25(0.35) | 90.14(7.25) |
| ALASSO | 0.54(0.07) | **0.59(0.06)** | **0.92(0.02)** | **0.07(0.05)** | **1.13(0.3)** | 6(2.74) |
| ENET | 0.71(0.07) | 0.44(0.08) | 0.91(0.03) | 0.09(0.05) | 1.33(0.4) | 54.96(5.22) |
| CB | **0.26(0.07)** | 0.56(0.05) | 0.86(0.04) | 0.09(0.05) | 1.22(0.25) | 52.09(2.08) |
| RSF | 0.97(0.01) | 0.06(0.01) | 0.73(0.06) | 0.2(0.04) | 2.3(0.4) | 666.44(37.42) |
| sRSF | 0.95(0.01) | 0.08(0.01) | 0.74(0.06) | 0.19(0.04) | 2.25(0.35) | 648.7(47.12) |
| BH | 0.97(0) | 0.06(0) | 0.52(0.07) | 0.49(0.09) | 4.41(1.1) | 21.33(5.19) |
| QV | 0.97(0) | 0.05(0) | 0.5(0.08) | 0.51(0.12) | 4.4(1.23) | 26.64(1.71) |
| CARS (MED) | 0.9(0.02) | 0.17(0.03) | 0.72(0.2) | 0.27(0.22) | 2.61(2.21) | **1.59(0.67)** |
| CARS (MSR) | 0.79(0.09) | 0.23(0.11) | 0.81(0.08) | 0.15(0.07) | 1.73(0.56) | 16.22(7.74) |

The entries are median(inter-quartile range). Well-performing methods have low FDR, high F1-score, high CI, low RMSE, and low computation time. The best median value for each metric is boldfaced. FDR: false discovery rate, CI: concordance index, RMSE: root mean squared error. Computation time was measured in seconds.

CoxBoost performed the best in controlling FDR, followed by ALASSO and LASSO. In contrast, BH and QV surprisingly had the worst performance under this metric. All embedded methods with the exception of RSF and sRSF had similarly good F1-scores, while the filter methods generally underperformed. The regularized methods LASSO, ALASSO, and ENET had very good CI and Brier score. CB also performed well in Brier score. ALASSO performed best in RMSE by a decent margin, and CARS with MED thresholding had the shortest computation times.

For quick reference, the bottom panel of Table 3 shows relative method rankings in this second setting of simulations.

### Real data analyses results

#### BLCA cohort analysis

The Kaplan-Meier estimators for BLCA males and females in Fig 4 show no significant differences in the survival functions between the two sexes (p=0.53) at the 5% level of significance. Survival probability dropped rapidly during the first 500 days following initial diagnosis, and tapered off after about 3,000 days, which suggests that many diagnoses may have occurred late in disease progression. Of 423 patients, 2, 133, 146, and 141 were at stages I through IV respectively (one patient’s stage was not recorded) at the time of initial surgical pathology or biopsy, and 208 of those procedures were within a year of the initial diagnoses. The median survival time was approximately 1,000 days for both genders (i.e., about half of the cohort experienced failure prior to this point). The presence of more males than females within the cohort leads to the 95% confidence intervals for the survival function being tighter for the males than for the females.

#### mRNA feature pairwise correlations

The histograms of APC between the 20,240 mRNA features prior to PFS and the 3,000 features following PFS are shown in Fig 5. Although a majority of feature pairs are not highly correlated, there still exist over 1.2 million feature pairs with APC exceeding 0.5 prior to PFS, as shown in the top panel. The distribution is somewhat more left-leaning following PFS, i.e. some feature pairs that had high APC were removed, as well as features with little to no association with the outcome.

#### mRNA feature selection

Summary statistics (median and inter-quartile range) of all feature selection metrics are provided in Table 5. Methods with higher ratios of true positives to selected mRNAs, higher Dice coefficient, and lesser computation times are better performing.

**Table 5.**
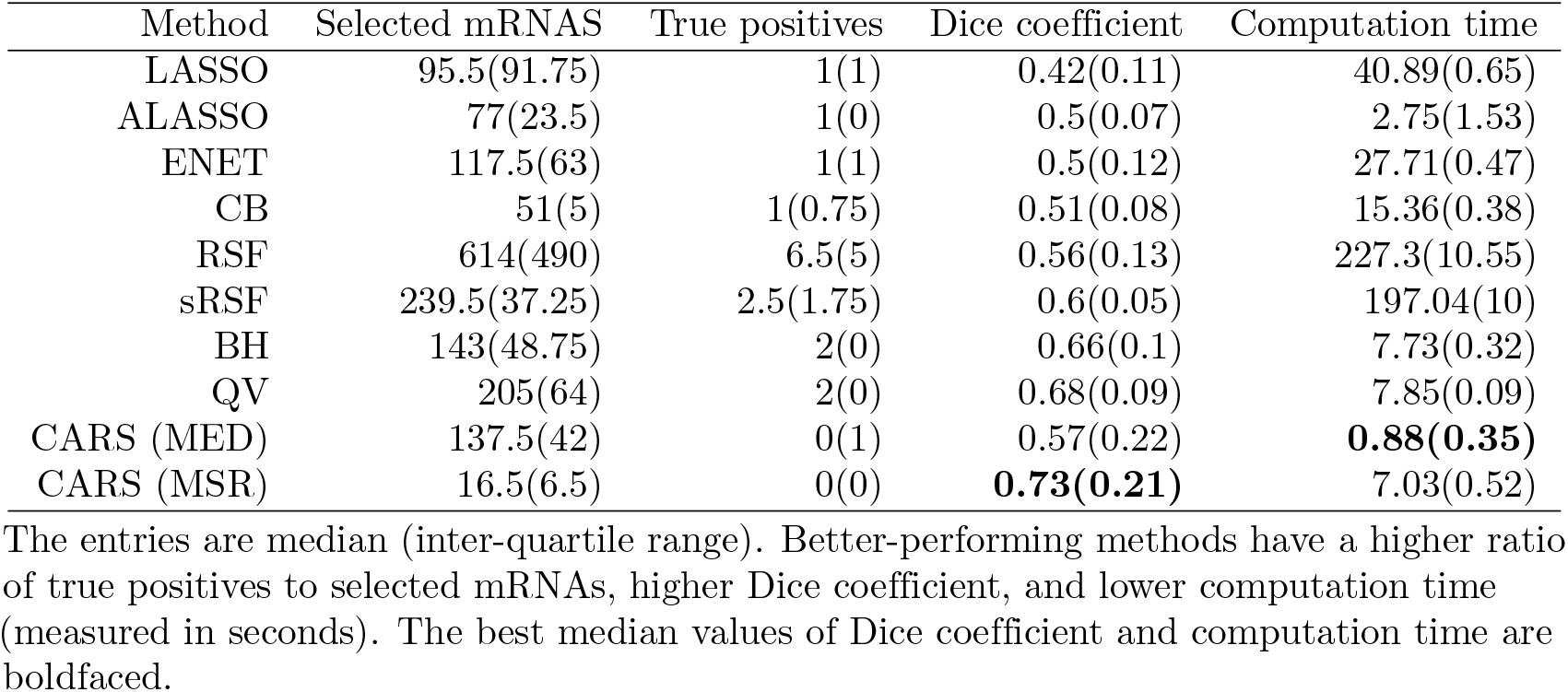
Feature selection and computation time metrics for the real data (TCGA-BLCA) analysis.

| Method | Selected mRNAs | True positives | Dice coefficient | Computation time |
| --- | --- | --- | --- | --- |
| LASSO | 95.5(91.75) | 1(1) | 0.42(0.11) | 40.89(0.65) |
| ALASSO | 77(23.5) | 1(0) | 0.5(0.07) | 2.75(1.53) |
| ENET | 117.5(63) | 1(1) | 0.5(0.12) | 27.71(0.47) |
| CB | 51(5) | 1(0.75) | 0.51(0.08) | 15.36(0.38) |
| RSF | 614(490) | 6.5(5) | 0.56(0.13) | 227.3(10.55) |
| sRSF | 239.5(37.25) | 2.5(1.75) | 0.6(0.05) | 197.04(10) |
| BH | 143(48.75) | 2(0) | 0.66(0.1) | 7.73(0.32) |
| QV | 205(64) | 2(0) | 0.68(0.09) | 7.85(0.09) |
| CARS (MED) | 137.5(42) | 0(1) | 0.57(0.22) | **0.88(0.35)** |
| CARS (MSR) | 16.5(6.5) | 0(0) | **0.73(0.21)** | 7.03(0.52) |

The entries are median (inter-quartile range). Better-performing methods have a higher ratio of true positives to selected mRNAs, higher Dice coefficient, and lower computation time (measured in seconds). The best median values of Dice coefficient and computation time are boldfaced.

Both RSF and sRSF selected the largest number of features, and RSF had the largest variation in this quantity over the ten outer folds. CB and CARS with MSR thresholding had consistently fewer selected features, and CB generally had at least one true positive. CARS with the MSR threshold also had the most consistent selected features (high Dice coefficient), followed by BH and QV. The LASSO had the worst Dice coefficient. CARS with the MED threshold had the least computation time.

The selection frequencies of the 30 mRNA features listed in Table 2 over the ten outer folds of the real data analysis are provided in S3 Table. The features DLEU1 and TOP3B were the most commonly selected by a majority of the methods.

### Prognostic modeling

Predictive performances based on the 10 folds of analysis on the real data are shown in Table 6. Methods with greater CI and lesser Brier score at both time points are better performing.

**Table 6.** Predictive metrics for the real data (TCGA-BLCA) analysis.

| Method | CI | Brier score (365 days) | Brier score (1,000 days) |
| --- | --- | --- | --- |
| LASSO | **0.59(0.06)** | 0.22(0.08) | 0.4(0.08) |
| ALASSO | 0.57(0.08) | 0.2(0.01) | 0.34(0.06) |
| ENET | **0.59(0.06)** | 0.23(0.11) | 0.36(0.13) |
| CB | 0.57(0.05) | 0.19(0.06) | 0.36(0.09) |
| RSF | 0.56(0.05) | **0.17(0.04)** | 0.28(0.01) |
| sRSF | **0.59(0.08)** | **0.17(0.04)** | 0.27(0.03) |
| BH | 0.57(0.07) | 0.23(0.06) | 0.36(0.03) |
| QV | 0.53(0.09) | 0.3(0.05) | 0.51(0.19) |
| CARS (MED) | 0.49(0.07) | 0.25(0.07) | 0.39(0.11) |
| CARS (MSR) | 0.56(0.16) | **0.17(0.05)** | **0.25(0.03)** |

The entries are median (inter-quartile range). Better-performing methods have higher CI and lower Brier score at both time points. The best median values of CI and Brier score are boldfaced. CI: concordance index.

Although the LASSO and ALASSO were the best performing in CI, sRSF was competitive in performance. CARS with MSR thresholding was the best in Brier score at both time points, and tied with both RSF methods at the 365 day horizon and slightly outperformed them at 1,000 days. Interestingly, although the parametric embedded methods (LASSO, ALASSO, ENET, CB) performed competitively in Brier score at 365 days, they greatly underperformed at 1,000 days.

#### Model calibration checks

We performed an additional heuristic check of the prediction performance by each method in the ten outer folds of the real data analysis. For each method and fold, observations were partitioned into quintiles based on predicted risk. We assessed model calibration by plotting predicted versus observed survival probabilities at the 1, 3, and 5-year clinical horizons. These plots are provided in S4 Fig, faceted by method and clinical horizon. The curves from the ten folds are colored and overlayed in each individual subplot. Well-calibrated models predict each group’s survival probability to be near their respective observed survival probabilities, shown visually by having survival curves near the solid black *y* = *x* line. Least squares regression lines are shown per method and horizon as dashed black lines, with their intercepts and slopes provided in each subplot.

Generally, the observed survival probabilities at the 1 year clinical horizon exceed 0.5, but they are more uniformly distributed at the 3 and 5-year horizons. Further examining the 1 year plots (the first and fourth columns), some methods have curves that are nearly vertical or even slightly downwards-sloping, suggesting that survival predictions may be volatile at this horizon, as there is greater deviation in predicted survival probabilities than in the observed ones. In the 3 and 5-year plots, many of the curves are biased above the *y* = *x* line, suggesting that the predicted survival probabilities are optimistic when the observed survival probability is about 0.5. This is largely the case with the parametric methods. The RSF and sRSF methods avoid this optimism, and have very polarized observed survival probabilities, indicative that the nonparametric forests have highly segmented the high and low risk groups. CARS with the MSR threshold has downward-sloping curves, which suggests that informative features may be missing from the final models. Overall, with few exceptions, most of the methods’ predictions are stable between the ten folds, shown by the colored curves being near each other in each subplot.

Note that the calibration we observe is largely dependent upon the underlying data characteristics. Methods’ performances change depending upon the data sparsity, strength of signals, predictor correlations, and censoring, many of which are unknown in the real data case. Further, it is important to examine calibration at multiple horizons, otherwise observed survival probabilities may be only high (as is the case in our 1 year plots) or only low.

## Discussion

### Comparison of CARS scores thresholding methods (MED, MSR)

For all replications of the first setting of simulations, we recorded the elbow point estimations (i.e. number of selected features) by MED and MSR to examine stability. Fig 9 shows the distributions of selected features by these thresholding methods by the three different levels of sparsity (proportion of true signals). As expected, when the number of truly important features increases, both thresholds tend to increase. The MSR threshold is consistently more stringent, and has a slightly tighter distribution.

**Fig 9.**
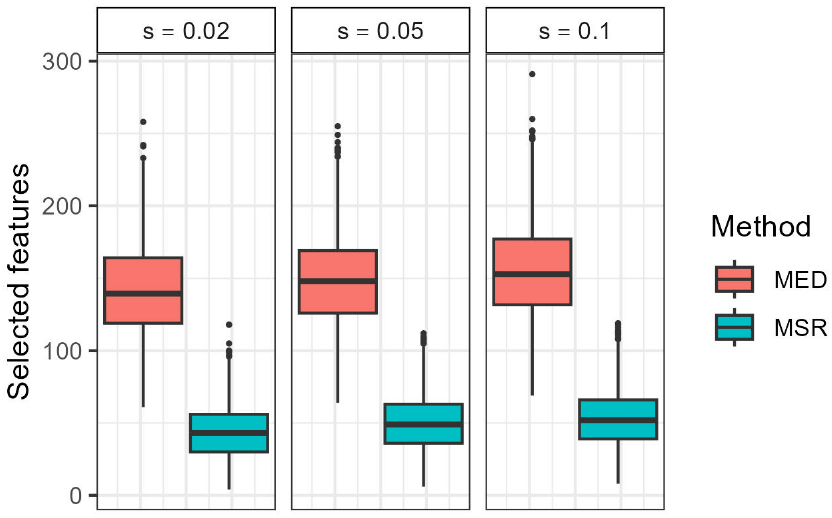
Distribution/boxplots of the number of selected features by MED and MSR thresholding methods for CARS scores. Plots are faceted by the three cases of sparsity (*s*) examined in the first setting of simulations.

### Excluded methods

Various other methods have been evaluated for this benchmark, but have been removed due to poor performance on our synthetic data.

#### Variance filter

The variance filter, which excelled in the benchmark study from Bommert et al. [5], is not applicable to our studies. In short, the variance filter gives each feature a score equal to its sample variance and selects the features with scores exceeding a specified threshold, which trivially makes it ineffective in our studies where features are standardized to uniform variance.

#### RSF with variable importance (VIMP)

A third variation of RSF from Ishwaran et al. [13] utilizes a different variable selection criterion called variable importance (VIMP). VIMP is the increase in prediction error if a particular variable had not been present within the data. Trivially, predictors with high VIMP are considered important. We examined the variable selection capability of RSF by choosing predictors with VIMP exceeding a threshold, but it performed worse than RSF by minimum depth with respect to both examined variable selection metrics.

#### Stepwise variable selection

A family of wrapper methods was also examined that utilized a screening procedure followed by stepwise variable selection. First, data dimensionality was reduced by selecting variables with the *n* − 2 lowest p-values from univariate Cox regression, or the variables with the lowest *n*_1_*/* log(*n*_1_) mean minimum depths after fitting a RSF, where 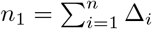, i.e. the number of uncensored observations within the data. Then a proportional hazards model was fit using iterative forward stepwise variable selection with either AIC or BIC as a model selection criterion. These methods were extremely computationally intensive and performed poorly in all metrics.

#### Bayesian additive regression trees (BART)

Much like the RSF, BART is a nonparametric tree ensemble method for estimating relationships between feature vectors and their associated outcomes. The model was introduced by Chipman et al. [66] in 2010 and adapted to right-censored survival data by Sparapani et al. [67] in 2016 with the corresponding R package `survbart`.

The general paradigm has inspiration from boosting methods where several weak learners (decision trees in this case) are fit sequentially, each successive one fitting to the residuals of the former so that the sum of these trees better captures the variations in the data. To prevent overfitting, regularized prior distributions of the model parameters (tree complexities and terminal node value estimates) are assumed, i.e. less complex decision trees are preferred. The corresponding posterior distributions are estimated using a Markov chain Monte Carlo (MCMC) algorithm, which repeatedly samples from the posterior [66].

We briefly examined the performance of BART in our predictive metrics (CI and RMSE) using the `survbart` package implementation. In the RMSE calculation, the estimated survival time was the median of the estimated survival function. In the first setting of our simulation study, BART was applied on 18 total synthetic datasets, one for each possible unique combination of our examined data characteristics.

The predictive performance of BART was promising and regularly outperformed the RSF/sRSF methods, but was extremely computationally taxing, and was therefore removed from the remainder of the study.

Park et al. [68] benchmarked the RSF and BART methods and some of their variations in terms of their predictive performances on synthetic linear regression data featuring missing values, high-dimensionality, and high predictor correlations. In their studies, they found that the DART, a variant of BART that utilizes Dirichlet prior distributions which encourage sparse models, outperformed the RSF in most of their predictive metrics, but suffered extreme computational costs.

The BART paradigm appears promising once improvements to the efficiency of the MCMC algorithm are made.

### Method performances

In simulation setting-I, the FDR-controlling methods BH and QV consistently had the lowest FDR and selected very few features. In contrast, in setting-II with weak to moderate signals, they had among the highest FDR, and in the real data analysis they typically selected about 140 and 210 features respectively, which was surprisingly large. These methods rely solely on univariate screening, which cannot capture gene–gene interactions or correlations. As a result, many features appear marginally correlated with the outcome and produce near-zero p-values, even though only a few true signals are present. This phenomenon is observed even after our PFS procedure reduces multicollinearity and dimensionality.

The nonparametric RSF and sRSF methods unsurprisingly performed better within the real data analysis than in the simulations, namely when performing prognostic modeling (prediction), as the proportional hazards model (1) may not hold for the real data. The poor performance of LASSO in terms of the Dice coefficient in the real data analysis is likely due to its instability in spaces with multi-collinear covariates.

Unsurprisingly, the embedded methods generally have greater predictive performance than the examined filter methods in our simulations. Although BH and QV excelled in controlling FDR in our first setting of simulations, they performed very poorly in the second setting and had a disproportionally large number of false positives. Further, none of the examined filters surpassed the parametric embedded methods (LASSO, ALASSO, ENET, CB) in F1-score. As an all-around performer, CARS with the MSR threshold is the best among the examined filter methods. This is unsurprising as CARS performed well in the filter method benchmark by Bommert et al. [5].

Considering both settings of simulations alone, we recommend the ALASSO and CB methods for general use due to their consistently good performances under both feature selection metrics and the three predictive metrics.

### Technical specifications

All studies were performed on the FASTER high-performance computing cluster maintained by Texas A&M University High Performance Research Computing. Every simulation was done on a single computing core with 248 gigabytes of RAM within an R [69] environment run on the Linux 8 operating system. Table 7 details the R packages used to implement the examined methods, and Table 8 outlines the existing hyperparameter tuning implementations for the examined methods in software. The specific usages of hyperparameter tuning and cross-validation in our studies are outlined in our descriptions of the examined methods.

**Table 7.**
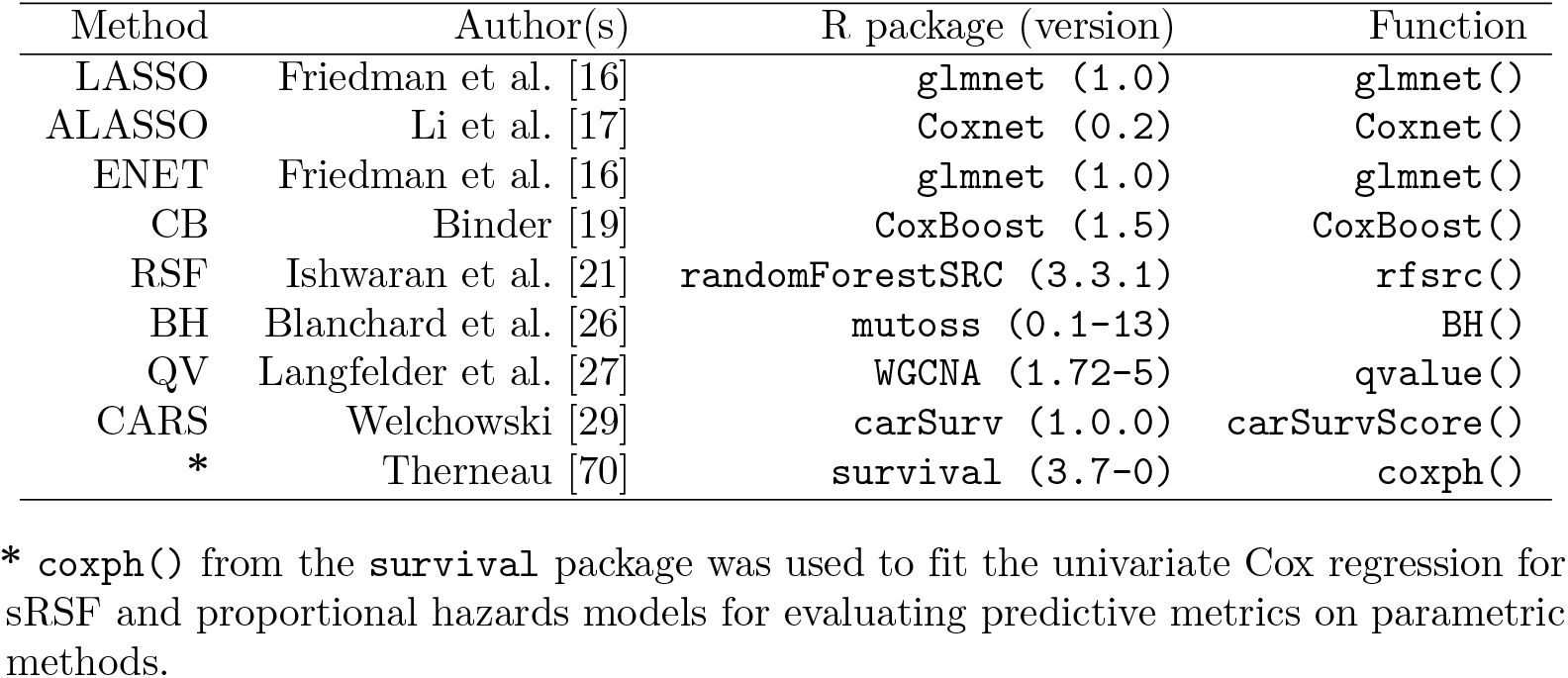
R package implementations for the examined methods.

**Table 8.** Existing hyperparameter tuning implementations for the examined methods.

| Method | Function | Tunable hyperparameter(s) | Search strategy | Search steps |
| --- | --- | --- | --- | --- |
| LASSO | `cv.glmnet()` | Regularization penalty $\lambda$ | Grid search | 100 |
| ALASSO | `Coxnet()` | Regularization penalty $\lambda$ | Grid search | 50 |
| ENET | `cv.glmnet()` | Regularization penalty $\lambda$ | Grid search | 100 |
| CB | `cv.CoxBoost()` | No. boosting steps $M$ | Grid search | 500 |
| *RSF | `tune.rfsrc()` | No. candidate splitting features<br>Minimum node size | Local search | $\approx 200$ |

Hyperparameter tuning implementations are provided by the methods’ respective R packages. The number of search steps may be adjustable in the function call, or vary depending on data dimensionality. Methods that perform grid search do so exhaustively. \* While other methods use cross-validation error estimates, RSF uses out-of-bag prediction error instead to guide the search.

## Conclusion

The tasks of biomarker identification and prognostic modeling are difficult on right-censored survival time and gene expression data, which features the problematic characteristics of right-censoring, sparse and high-dimensional features, and high correlations between features. Using the programming language R, we have evaluated a diverse selection of nine prominent methods for performing the aforementioned tasks in a simulation study to determine the best performers under various metrics and data characteristics. We have also performed analyses on a cohort of real cancer patients (TCGA-BLCA) to illustrate how the examined methods can be applied to real data, and designed a complementary set of simulations that closely mimic the characteristics of this cohort, allowing us to assess method performance under conditions where the true outcomes were known. The main results of the study are summarized below.

Regularized methods, namely ALASSO and CoxBoost were the most effective at developing prognostic models with well-chosen feature subsets, and we recommend them for general use. The non-parametric Random Survival Forest was almost universally improved by reducing data dimensionality with a filter method as an initial step. Of the two classical FDR-controlling methods we considered when selecting features with the univariate Cox regression, both the Benjamini-Hoschberg and q-value procedures show varying degree of performance depending on the correlation among the features and strength of true signals. Therefore, we advise against exclusive use of these univariate feature selection methods. In contrast, the performance of the CARS filter was far more consistent in our studies and we recommend its use when reduction of data dimensionality is needed. Of the two data-dependent thresholding techniques considered when performing CARS, our novel ad-hoc approach for estimating the selection threshold (MSR) generally outperformed the more commonly used technique (MED) for this purpose. Among the three regularized methods examined (LASSO, ALASSO, elastic net), we found that the ALASSO was consistently the best performer in our simulation studies.

## Acknowledgments

Simulations were conducted with computing resources provided by Texas A&M High Performance Research Computing. The results shown here are in part based upon data generated by the TCGA Research Network: https://www.cancer.gov/tcga. All plots were made using the R packages `ggplot2` [71] or `survminer` [72]. The data-flow diagrams (Figs 2, 3, and 6) were created in Lucid (`lucid.co`). The authors thank the reviewers for their constructive comments, which helped to significantly improve the quality of the article.

## Availability of data and materials

All R code to perform the benchmark and recreate our figures and tables are available in our GitHub repository https://github.com/WesLFletch/censored-data-regression-benchmark, commit hash cb1e548c54fd142e7a11c5951608366accc91478. TCGA-BLCA cohort data is publicly accessible from the Xena browser https://xenabrowser.net/datapages/ [34], or using the R package `UCSCXenaTools`. A timestamped copy of the cohort data and accompanying code are available in our repository.

## Supporting information

**S1 Appendix. Additional studies**. Five additional studies that were performed are presented: a study on how the choice of tuning parameters affects the predictive performance of RSF; a comparison between held-out (used in our studies) and out-of-bag predictive metrics for evaluating RSF; an examination of how our PFS procedure affects downstream prognostic modeling performance by the embedded methods on the real data; a simulation study that examines the relationship between the signal-to-noise ratios of true signals and their retention rate by PFS (using the CARS filter); and an analysis of the stability of PFS on the real BLCA data.

**S2 Table. Table of median metric values and inter-quartile ranges by all examined methods in the first setting of simulations**. Table rows are partitioned by method and metric. Columns are partitioned data characteristics (*α, s, γ*). Higher values of CI and F1-score, and lower values of Brier score, RMSE and FDR and computation time indicate better performance. CI: Concordance Index, RMSE: Root Mean Squared Error, FDR: False Discovery Rate.

**S3 Table. Real data analysis mRNA feature selection frequencies by the examined methods over the ten outer folds**. The 30 mRNA features likely associated with differences in survival time listen in Table 2 are listed over the rows, and the examined methods are provided in the columns. Each cell corresponds to how many of the ten outer folds the examined method selected the respective mRNA feature.

**S4 Fig. Calibration plots for the real data analysis**. The calibration curves of predicted-on-observed survival probabilities are faceted by method and clinical horizon. The curves for each of the 10 folds are colored and overlayed within each subplot. Well-calibrated methods have curves near the *y* = *x* line, shown in solid black. Least squares regression lines are shown per method and horizon as dashed black lines, with their intercepts and slopes provided in each subplot.

