## Supplementary figures and images for "Benchmark of biomarker identification and prognostic modeling methods on diverse censored data"

### Supporting Information 4: Figure

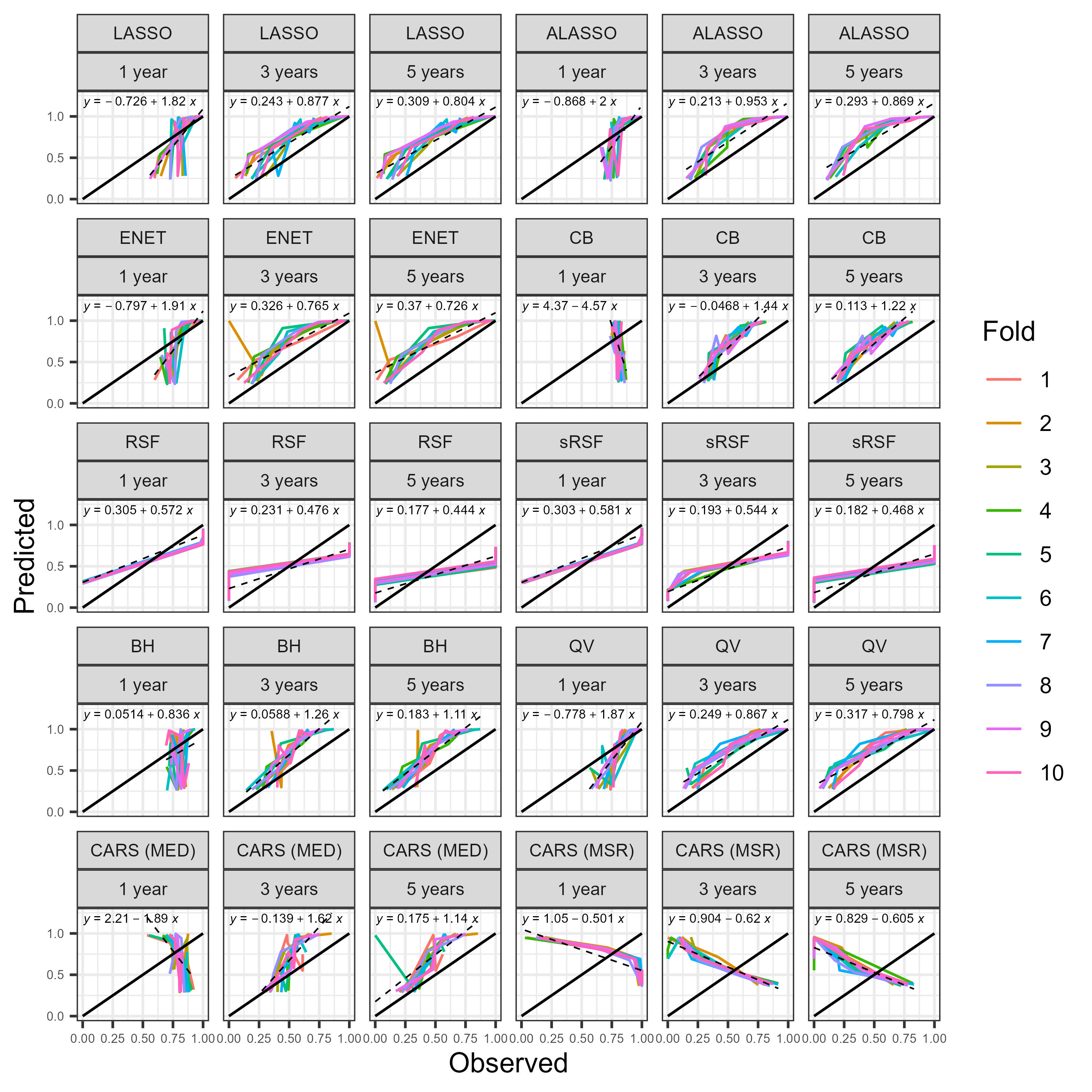
