## Supporting Information 2: Table for "Benchmark of biomarker identification and prognostic modeling methods on diverse censored data"

| | | $\alpha = 0$ | | | | | | | | | $\alpha = 0.5$ | | | | | | | | |
|---|---|---|---|---|---|---|---|---|---|---|---|---|---|---|---|---|---|---|---|
| | | $s = 0.02$ | | | $s = 0.05$ | | | $s = 0.1$ | | | $s = 0.02$ | | | $s = 0.05$ | | | $s = 0.1$ | | |
| | | $\gamma = 0.5$ | $\gamma = 1$ | $\gamma = 2$ | $\gamma = 0.5$ | $\gamma = 1$ | $\gamma = 2$ | $\gamma = 0.5$ | $\gamma = 1$ | $\gamma = 2$ | $\gamma = 0.5$ | $\gamma = 1$ | $\gamma = 2$ | $\gamma = 0.5$ | $\gamma = 1$ | $\gamma = 2$ | $\gamma = 0.5$ | $\gamma = 1$ | $\gamma = 2$ |
| FDR | LASSO | 0.75(0.12) | 0.73(0.1) | 0.71(0.14) | 0.75(0.1) | 0.74(0.09) | 0.74(0.08) | 0.71(0.11) | 0.69(0.11) | 0.7(0.11) | 0.78(0.13) | 0.76(0.07) | 0.73(0.11) | 0.76(0.11) | 0.74(0.11) | 0.73(0.07) | 0.72(0.11) | 0.71(0.1) | 0.71(0.1) |
| | ALASSO | 0.68(0.07) | 0.7(0.07) | 0.74(0.04) | 0.57(0.08) | 0.54(0.09) | 0.53(0.09) | 0.52(0.11) | 0.5(0.1) | 0.5(0.1) | 0.72(0.1) | 0.69(0.06) | 0.72(0.04) | 0.61(0.12) | 0.55(0.09) | 0.52(0.09) | 0.55(0.14) | 0.5(0.12) | 0.5(0.13) |
| | ENET | 0.78(0.11) | 0.78(0.08) | 0.76(0.07) | 0.79(0.1) | 0.78(0.09) | 0.77(0.09) | 0.76(0.13) | 0.79(0.12) | 0.79(0.12) | 0.82(0.12) | 0.79(0.08) | 0.78(0.06) | 0.8(0.11) | 0.79(0.11) | 0.79(0.08) | 0.79(0.12) | 0.78(0.1) | 0.78(0.1) |
| | CB | 0.48(0.12) | 0.26(0.14) | 0.19(0.13) | 0.53(0.1) | 0.48(0.11) | 0.46(0.1) | 0.55(0.1) | 0.54(0.11) | 0.53(0.1) | 0.56(0.15) | 0.29(0.17) | 0.2(0.17) | 0.55(0.15) | 0.48(0.14) | 0.42(0.14) | 0.55(0.14) | 0.5(0.14) | 0.5(0.14) |
| | RSF | 0.97(0.01) | 0.97(0.01) | 0.97(0.01) | 0.94(0.01) | 0.94(0.01) | 0.94(0.01) | 0.89(0.02) | 0.89(0.02) | 0.89(0.02) | 0.97(0.01) | 0.96(0.01) | 0.96(0.01) | 0.93(0.02) | 0.93(0.02) | 0.93(0.02) | 0.89(0.03) | 0.89(0.02) | 0.88(0.02) |
| | sRSF | 0.69(0.08) | 0.65(0.09) | 0.63(0.09) | 0.6(0.09) | 0.57(0.1) | 0.56(0.09) | 0.53(0.1) | 0.52(0.09) | 0.51(0.1) | 0.67(0.29) | 0.58(0.44) | 0.56(0.42) | 0.56(0.29) | 0.52(0.31) | 0.51(0.28) | 0.58(0.23) | 0.55(0.23) | 0.55(0.26) |
| | BH | **0(0)** | **0(0)** | **0(0)** | **0(0)** | **0(0)** | **0(0)** | **0(0)** | **0(0)** | **0(0)** | **0(0)** | **0(0)** | **0(0)** | **0(0)** | **0(0)** | **0(0)** | **0(0)** | **0(0)** | **0(0)** |
| | QV | **0(0)** | **0(0)** | **0(0)** | **0(0)** | **0(0)** | **0(0)** | **0(0)** | **0(0)** | **0(0)** | **0(0)** | **0(0)** | **0(0)** | **0(0)** | **0(0)** | **0(0)** | **0(0)** | **0(0)** | **0(0)** |
| | CARS (MED) | 0.92(0.03) | 0.9(0.03) | 0.88(0.03) | 0.86(0.03) | 0.84(0.04) | 0.83(0.04) | 0.8(0.04) | 0.79(0.04) | 0.78(0.04) | 0.93(0.02) | 0.9(0.03) | 0.88(0.03) | 0.87(0.03) | 0.84(0.04) | 0.83(0.03) | 0.81(0.04) | 0.79(0.04) | 0.78(0.04) |
| | CARS (MSR) | 0.84(0.07) | 0.76(0.1) | 0.72(0.12) | 0.77(0.08) | 0.73(0.09) | 0.71(0.11) | 0.72(0.08) | 0.69(0.08) | 0.69(0.09) | 0.88(0.06) | 0.79(0.09) | 0.71(0.14) | 0.81(0.08) | 0.73(0.1) | 0.7(0.09) | 0.75(0.09) | 0.71(0.09) | 0.69(0.09) |
| F1-score | LASSO | 0.39(0.14) | 0.42(0.13) | 0.45(0.17) | 0.35(0.09) | 0.4(0.09) | 0.4(0.08) | **0.33(0.07)** | **0.34(0.06)** | **0.35(0.06)** | 0.33(0.14) | 0.38(0.09) | 0.42(0.13) | 0.32(0.09) | 0.38(0.11) | 0.4(0.08) | **0.31(0.07)** | **0.33(0.06)** | **0.34(0.06)** |
| | ALASSO | 0.47(0.08) | 0.47(0.08) | 0.41(0.05) | **0.45(0.09)** | **0.51(0.09)** | **0.52(0.08)** | 0.3(0.08) | 0.31(0.09) | 0.32(0.08) | 0.39(0.11) | 0.47(0.07) | 0.44(0.05) | **0.4(0.09)** | **0.49(0.08)** | **0.53(0.07)** | 0.28(0.09) | 0.32(0.09) | 0.32(0.1) |
| | ENET | 0.35(0.14) | 0.36(0.11) | 0.38(0.09) | 0.33(0.11) | 0.35(0.11) | 0.36(0.1) | 0.31(0.05) | 0.31(0.05) | 0.31(0.05) | 0.29(0.14) | 0.34(0.1) | 0.36(0.08) | 0.3(0.11) | 0.33(0.13) | 0.34(0.1) | 0.29(0.05) | 0.31(0.06) | 0.32(0.05) |
| | CB | **0.61(0.11)** | **0.8(0.11)** | **0.85(0.1)** | 0.39(0.08) | 0.43(0.08) | 0.44(0.07) | 0.25(0.05) | 0.26(0.05) | 0.26(0.05) | **0.47(0.14)** | **0.68(0.13)** | **0.75(0.11)** | 0.31(0.09) | 0.35(0.09) | 0.37(0.1) | 0.19(0.06) | 0.21(0.06) | 0.21(0.05) |
| | RSF | 0.05(0.02) | 0.06(0.02) | 0.06(0.02) | 0.1(0.02) | 0.1(0.02) | 0.1(0.02) | 0.17(0.03) | 0.17(0.03) | 0.17(0.03) | 0.06(0.02) | 0.07(0.02) | 0.07(0.03) | 0.11(0.03) | 0.12(0.03) | 0.12(0.03) | 0.17(0.04) | 0.17(0.04) | 0.17(0.03) |
| | sRSF | 0.41(0.1) | 0.46(0.09) | 0.48(0.09) | 0.36(0.08) | 0.39(0.08) | 0.38(0.08) | 0.27(0.06) | 0.28(0.06) | 0.26(0.05) | 0.29(0.15) | 0.38(0.22) | 0.4(0.2) | 0.23(0.09) | 0.25(0.1) | 0.25(0.1) | 0.16(0.14) | 0.17(0.12) | 0.16(0.13) |
| | BH | 0.18(0.2) | 0.31(0.22) | 0.33(0.17) | 0(0.04) | 0(0.04) | 0(0.04) | 0(0) | 0(0) | 0(0) | 0(0) | 0(0) | 0(0.1) | 0(0) | 0(0) | 0(0) | 0(0) | 0(0) | 0(0) |
| | QV | 0.18(0.22) | 0.32(0.22) | 0.33(0.18) | 0(0.04) | 0(0.04) | 0(0.04) | 0(0) | 0(0) | 0(0) | 0(0) | 0(0.04) | 0(0.04) | 0(0) | 0(0) | 0(0) | 0(0) | 0(0) | 0(0) |
| | CARS (MED) | 0.15(0.04) | 0.18(0.05) | 0.2(0.06) | 0.22(0.04) | 0.24(0.05) | 0.25(0.05) | 0.24(0.04) | 0.26(0.04) | 0.27(0.04) | 0.12(0.04) | 0.17(0.05) | 0.2(0.05) | 0.19(0.05) | 0.23(0.05) | 0.25(0.04) | 0.23(0.03) | 0.26(0.04) | 0.27(0.05) |
| | CARS (MSR) | 0.23(0.08) | 0.32(0.1) | 0.37(0.1) | 0.23(0.07) | 0.27(0.08) | 0.28(0.07) | 0.19(0.07) | 0.21(0.06) | 0.21(0.07) | 0.17(0.07) | 0.29(0.1) | 0.38(0.12) | 0.19(0.08) | 0.25(0.08) | 0.29(0.08) | 0.18(0.07) | 0.2(0.07) | 0.22(0.07) |
| CI | LASSO | **0.75(0.05)** | 0.88(0.03) | **0.95(0.01)** | 0.66(0.06) | **0.72(0.06)** | **0.77(0.08)** | 0.61(0.06) | 0.63(0.06) | 0.66(0.08) | 0.66(0.06) | **0.83(0.04)** | **0.92(0.02)** | **0.63(0.06)** | **0.69(0.05)** | **0.74(0.05)** | 0.6(0.06) | 0.62(0.06) | 0.64(0.07) |
| | ALASSO | **0.75(0.06)** | **0.89(0.03)** | **0.95(0.01)** | 0.65(0.06) | 0.7(0.06) | 0.73(0.07) | 0.6(0.05) | 0.61(0.06) | 0.63(0.07) | **0.67(0.06)** | **0.83(0.05)** | **0.92(0.02)** | **0.63(0.05)** | 0.68(0.05) | 0.71(0.06) | 0.59(0.06) | 0.61(0.05) | 0.62(0.06) |
| | ENET | 0.74(0.06) | 0.86(0.03) | 0.92(0.02) | **0.67(0.05)** | 0.71(0.06) | 0.75(0.07) | **0.63(0.06)** | **0.66(0.06)** | **0.68(0.06)** | 0.65(0.06) | 0.8(0.05) | 0.88(0.03) | **0.63(0.05)** | **0.69(0.05)** | 0.72(0.05) | **0.62(0.06)** | **0.64(0.05)** | **0.66(0.05)** |
| | CB | 0.71(0.06) | 0.78(0.05) | 0.81(0.04) | 0.63(0.05) | 0.64(0.05) | 0.67(0.06) | 0.59(0.04) | 0.6(0.05) | 0.61(0.06) | 0.64(0.06) | 0.71(0.05) | 0.74(0.05) | 0.59(0.05) | 0.61(0.05) | 0.62(0.06) | 0.56(0.05) | 0.57(0.06) | 0.58(0.06) |
| | RSF | 0.52(0.06) | 0.52(0.05) | 0.53(0.05) | 0.52(0.05) | 0.52(0.05) | 0.52(0.05) | 0.51(0.05) | 0.52(0.05) | 0.51(0.06) | 0.51(0.06) | 0.52(0.05) | 0.52(0.05) | 0.51(0.05) | 0.51(0.04) | 0.51(0.05) | 0.51(0.05) | 0.5(0.05) | 0.51(0.06) |
| | sRSF | 0.57(0.05) | 0.59(0.05) | 0.59(0.05) | 0.55(0.05) | 0.56(0.05) | 0.56(0.05) | 0.55(0.06) | 0.55(0.05) | 0.55(0.06) | 0.54(0.06) | 0.57(0.08) | 0.58(0.08) | 0.53(0.06) | 0.54(0.06) | 0.54(0.06) | 0.52(0.06) | 0.52(0.05) | 0.52(0.06) |
| | BH | 0.58(0.08) | 0.63(0.08) | 0.65(0.07) | 0.5(0.05) | 0.5(0.05) | 0.5(0.04) | 0.5(0) | 0.5(0) | 0.5(0) | 0.5(0) | 0.5(0) | 0.5(0) | 0.5(0) | 0.5(0) | 0.5(0) | 0.5(0) | 0.5(0) | 0.5(0) |
| | QV | 0.59(0.09) | 0.64(0.08) | 0.65(0.07) | 0.5(0.05) | 0.5(0.05) | 0.5(0.04) | 0.5(0) | 0.5(0) | 0.5(0) | 0.5(0) | 0.5(0.01) | 0.5(0.02) | 0.5(0) | 0.5(0) | 0.5(0) | 0.5(0) | 0.5(0) | 0.5(0) |
| | CARS (MED) | 0.58(0.07) | 0.66(0.08) | 0.7(0.09) | 0.56(0.06) | 0.58(0.07) | 0.59(0.07) | 0.54(0.05) | 0.55(0.05) | 0.55(0.08) | 0.55(0.08) | 0.63(0.09) | 0.7(0.1) | 0.55(0.06) | 0.58(0.07) | 0.59(0.07) | 0.54(0.06) | 0.56(0.05) | 0.55(0.07) |
| | CARS (MSR) | 0.63(0.06) | 0.69(0.07) | 0.72(0.06) | 0.59(0.05) | 0.61(0.05) | 0.62(0.05) | 0.57(0.05) | 0.57(0.05) | 0.58(0.07) | 0.58(0.06) | 0.66(0.08) | 0.71(0.08) | 0.58(0.06) | 0.6(0.06) | 0.62(0.05) | 0.56(0.05) | 0.57(0.05) | 0.58(0.05) |
| Brier score | LASSO | 0.23(0.07) | **0.11(0.05)** | **0.08(0.03)** | 0.35(0.14) | 0.32(0.16) | 0.36(0.19) | 0.38(0.14) | 0.39(0.16) | 0.41(0.28) | 0.29(0.09) | 0.17(0.06) | **0.08(0.04)** | 0.35(0.11) | 0.32(0.12) | 0.33(0.16) | 0.38(0.13) | 0.39(0.14) | 0.4(0.15) |
| | ALASSO | 0.21(0.06) | **0.11(0.04)** | **0.08(0.03)** | 0.27(0.05) | **0.23(0.06)** | **0.22(0.04)** | 0.29(0.05) | 0.28(0.05) | 0.25(0.04) | 0.27(0.08) | **0.15(0.05)** | **0.08(0.04)** | 0.29(0.05) | **0.25(0.06)** | **0.23(0.05)** | 0.29(0.05) | 0.29(0.06) | 0.27(0.04) |
| | ENET | 0.25(0.08) | 0.13(0.06) | 0.1(0.04) | 0.43(0.16) | 0.37(0.18) | 0.45(0.17) | 0.43(0.16) | 0.46(0.14) | 0.57(0.21) | 0.3(0.12) | 0.19(0.08) | 0.09(0.04) | 0.41(0.17) | 0.41(0.2) | 0.46(0.16) | 0.46(0.16) | 0.44(0.12) | 0.47(0.12) |
| | CB | **0.19(0.05)** | **0.11(0.05)** | 0.09(0.04) | 0.27(0.05) | 0.24(0.05) | 0.23(0.04) | 0.29(0.04) | 0.28(0.04) | 0.26(0.04) | **0.24(0.05)** | 0.17(0.04) | 0.15(0.04) | 0.27(0.04) | **0.25(0.04)** | 0.25(0.04) | 0.29(0.04) | 0.28(0.04) | 0.27(0.04) |
| | RSF | 0.26(0.02) | 0.26(0.02) | 0.26(0.02) | 0.26(0.02) | 0.26(0.02) | 0.26(0.02) | 0.26(0.02) | 0.26(0.02) | 0.26(0.02) | 0.28(0.03) | 0.28(0.03) | 0.28(0.03) | 0.28(0.03) | 0.28(0.03) | 0.28(0.03) | 0.28(0.03) | 0.28(0.03) | 0.28(0.03) |
| | sRSF | 0.25(0.02) | 0.24(0.02) | 0.24(0.02) | **0.25(0.02)** | 0.25(0.02) | 0.25(0.02) | 0.26(0.02) | 0.26(0.02) | 0.26(0.02) | 0.26(0.03) | 0.25(0.03) | 0.25(0.03) | 0.26(0.03) | 0.26(0.03) | 0.26(0.03) | 0.27(0.02) | 0.27(0.03) | 0.27(0.03) |
| | BH | 0.25(0.02) | 0.23(0.03) | 0.22(0.03) | **0.25(0.01)** | 0.25(0.01) | 0.25(0.01) | **0.25(0.01)** | **0.25(0.01)** | **0.24(0.02)** | 0.25(0.01) | 0.25(0) | 0.25(0) | **0.25(0)** | **0.25(0)** | 0.25(0) | **0.25(0)** | **0.25(0)** | **0.25(0)** |
| | QV | 0.25(0.02) | 0.23(0.03) | 0.22(0.03) | **0.25(0.01)** | 0.25(0.01) | 0.25(0.01) | **0.25(0.01)** | **0.25(0.01)** | **0.24(0.02)** | 0.25(0.01) | 0.25(0.01) | 0.25(0) | **0.25(0)** | **0.25(0)** | 0.25(0) | **0.25(0.01)** | **0.25(0)** | **0.25(0)** |
| | CARS (MED) | 0.39(0.11) | 0.31(0.11) | 0.25(0.1) | 0.41(0.08) | 0.38(0.09) | 0.37(0.11) | 0.42(0.09) | 0.41(0.09) | 0.4(0.14) | 0.41(0.1) | 0.34(0.12) | 0.26(0.1) | 0.41(0.09) | 0.38(0.1) | 0.36(0.12) | 0.43(0.1) | 0.41(0.1) | 0.4(0.11) |
| | CARS (MSR) | 0.26(0.05) | 0.22(0.05) | 0.19(0.04) | 0.28(0.05) | 0.27(0.05) | 0.25(0.03) | 0.29(0.06) | 0.29(0.05) | 0.27(0.04) | 0.28(0.05) | 0.23(0.05) | 0.2(0.06) | 0.29(0.05) | 0.27(0.05) | 0.26(0.04) | 0.3(0.05) | 0.29(0.05) | 0.28(0.04) |
| RMSE | LASSO | 2.02(0.3) | 2.47(0.57) | **4.5(1.42)** | 3.61(0.56) | 6.62(1.07) | 18.36(7.69) | 5.33(0.7) | 11.11(2.86) | 36.64(20.98) | 1.93(0.31) | 2.18(0.45) | 3.2(0.89) | 2.82(0.31) | 4.78(0.85) | 10.46(1.87) | 3.9(0.59) | 7.3(0.99) | 19.95(9.36) |
| | ALASSO | **1.94(0.29)** | 2.36(0.4) | 4.53(1.2) | **3.47(0.41)** | **6.12(0.83)** | 20.79(5.02) | 5.22(0.66) | 12.1(2.12) | 43.21(8.64) | 1.88(0.27) | **2.02(0.36)** | **3.19(0.6)** | **2.76(0.33)** | **4.48(0.57)** | 10.33(1.8) | 3.85(0.38) | 7.25(0.83) | 23.72(5.35) |
| | ENET | 2.11(0.34) | 2.63(0.63) | 4.97(1.3) | 3.86(0.89) | 6.87(1.03) | 16.1(7.07) | 5.43(1.01) | 10.76(1.88) | 23.53(13.92) | 1.97(0.27) | 2.33(0.52) | 3.33(0.88) | 2.99(0.67) | 5.1(1.1) | 10.41(1.63) | 4.12(1.2) | **7.14(1.15)** | 15.34(4.43) |
| | CB | 1.96(0.31) | **2.34(0.63)** | 5.14(1.35) | 3.62(0.45) | 6.66(0.86) | 22.19(4.64) | 5.31(0.56) | 12.57(2.2) | 43.58(9.16) | **1.86(0.28)** | 2.43(0.37) | 4.05(0.74) | 2.84(0.3) | 4.95(0.56) | 11.74(2.13) | 3.94(0.4) | 7.46(0.75) | 24.36(5.71) |
| | RSF | 2.63(0.25) | 4.77(0.49) | 9.25(0.88) | 3.91(0.39) | 7.43(0.73) | 14.73(1.55) | 5.37(0.56) | 10.45(1.14) | 20.81(2.25) | 2.24(0.24) | 3.73(0.34) | 7.04(0.68) | 3.14(0.31) | 5.72(0.55) | 11.17(1.18) | 4.18(0.45) | 7.97(0.87) | 15.72(1.62) |
| | sRSF | 2.56(0.24) | 4.52(0.5) | 8.76(0.89) | 3.82(0.37) | 7.26(0.67) | **14.27(1.44)** | 5.31(0.56) | 10.28(1.1) | **20.48(2.37)** | 2.11(0.23) | 3.42(0.41) | 6.36(0.68) | 2.94(0.32) | 5.36(0.54) | 10.47(1.1) | 3.97(0.45) | 7.62(0.88) | 15.03(1.63) |
| | BH | 2.49(0.25) | 4.28(0.48) | 8.5(1.12) | 3.79(0.4) | 7.22(0.69) | 15.87(13.78) | **5.2(0.52)** | **10.17(1.2)** | 56.61(11.92) | 2.04(0.2) | 3.4(0.29) | 6.41(0.57) | 2.81(0.27) | 5.17(0.47) | **10.12(1.03)** | **3.76(0.37)** | **7.14(0.72)** | **14.24(1.81)** |
| | QV | 2.49(0.25) | 4.27(0.49) | 8.5(1.13) | 3.79(0.4) | 7.22(0.69) | 15.95(13.52) | **5.2(0.52)** | 10.19(1.2) | 56.6(11.94) | 2.05(0.21) | 3.42(0.3) | 6.43(0.57) | 2.82(0.28) | 5.18(0.47) | **10.12(1)** | 3.79(0.45) | **7.14(0.69)** | **14.28(2.04)** |
| | CARS (MED) | 3.28(0.9) | 4.74(1.29) | 9.71(2.61) | 5(1.03) | 9.19(2.71) | 26.38(6.47) | 7.05(1.8) | 16.75(4.89) | 44.2(11.92) | 2.85(0.82) | 3.75(1.03) | 5.81(1.5) | 3.75(0.91) | 6.35(1.37) | 14.78(3.94) | 5(1.14) | 9.62(2.39) | 27.25(6.86) |
| | CARS (MSR) | 2.49(0.35) | 3.91(0.67) | 8.12(1.22) | 3.92(0.48) | 7.3(0.83) | 24.9(5.05) | 5.5(0.75) | 13.27(2.41) | 46.69(9.4) | 2.11(0.29) | 3.04(0.49) | 5.11(0.83) | 2.96(0.34) | 5.21(0.54) | 12.31(2.47) | 4.06(0.45) | 7.65(0.77) | 26.16(6.08) |
| Computation time | LASSO | 36.43(3.85) | 39.31(4.14) | 41.71(4.79) | 36.24(4.12) | 37.36(4.35) | 37.8(4.61) | 36.4(3.54) | 37.56(3.93) | 38.18(4.13) | 35.78(6.02) | 41.44(7.45) | 48.77(8.46) | 35.97(5.23) | 38.01(6) | 38.71(6.14) | 36.6(6.38) | 38.07(7.97) | 37.58(7.87) |
| | ALASSO | 1.04(0.35) | 1.08(0.24) | 1.04(0.31) | 1(0.42) | 1(0.57) | 1.03(0.49) | 1.06(0.51) | 1.06(0.57) | 1.07(0.49) | 1.72(0.38) | 1.6(0.41) | 1.53(0.47) | 1.89(0.5) | 1.86(0.55) | 1.87(0.62) | 2.03(0.49) | 2.07(0.59) | 2.06(0.54) |
| | ENET | 18.98(1.75) | 18.53(1.54) | 18.36(1.88) | 19.6(1.87) | 19.74(1.85) | 20(2.03) | 20.46(2.67) | 21.16(2.83) | 21.73(3) | 21.04(3.07) | 21.77(2.87) | 22.47(3.2) | 22.27(2.68) | 23.02(2.6) | 23.46(2.79) | 23.55(3.79) | 23.69(4.09) | 23.74(5.02) |
| | CB | 4.92(0.27) | 5(0.09) | 5.06(0.08) | 4.95(0.11) | 5.05(0.1) | 5.09(0.08) | 5.01(0.09) | 5.08(0.09) | 5.1(0.09) | 4.81(0.15) | 4.95(0.11) | 5.05(0.09) | 4.9(0.13) | 5.03(0.1) | 5.09(0.08) | 4.97(0.12) | 5.05(0.12) | 5.1(0.14) |
| | RSF | 365.22(18.93) | 377.24(18.68) | 383.77(15.12) | 373.25(16.48) | 387.79(13.15) | 392.33(14.89) | 382.58(15.19) | 391.05(14.48) | 393.84(13.25) | 200.43(28.26) | 205.34(28.27) | 210.76(29.17) | 208.93(28.55) | 215.64(35.35) | 219.3(35.22) | 209.91(33.61) | 214.56(38.76) | 213.94(47.54) |
| | sRSF | 56.97(9.94) | 59.1(9.51) | 59.3(9.21) | 62.36(9.13) | 64.58(9.95) | 60.89(9.31) | 66.02(11.2) | 66.21(11.48) | 55.47(10.18) | 26.46(34.86) | 27.4(40.21) | 28.27(39.85) | 25.26(26.81) | 26.83(23.64) | 26.36(29.09) | 32.87(49.15) | 30.62(43.27) | 25.77(37.62) |
| | BH | 4.75(0.55) | 4.76(0.35) | 4.85(0.29) | 4.74(0.25) | 4.88(0.2) | 4.84(0.24) | 4.79(0.26) | 4.85(0.21) | 4.83(0.23) | 4.77(0.19) | 4.75(0.2) | 4.94(0.17) | 4.79(0.19) | 4.86(0.21) | 4.91(0.19) | 4.77(0.37) | 4.87(0.89) | 4.82(2.32) |
| | QV | 4.75(0.47) | 4.76(0.34) | 4.83(0.28) | 4.72(0.24) | 4.86(0.24) | 4.81(0.24) | 4.8(0.24) | 4.83(0.19) | 4.83(0.23) | 4.75(0.17) | 4.74(0.16) | 4.92(0.19) | 4.77(0.17) | 4.84(0.2) | 4.9(0.18) | 4.75(0.53) | 4.88(1.23) | 4.82(1.43) |
| | CARS (MED) | **0.26(0.38)** | **0.25(0.02)** | **0.25(0.02)** | **0.24(0.02)** | **0.25(0.02)** | **0.25(0.02)** | **0.25(0.02)** | **0.25(0.02)** | **0.25(0.02)** | **0.26(0.03)** | **0.25(0.02)** | **0.25(0.02)** | **0.25(0.02)** | **0.25(0.03)** | **0.26(0.03)** | **0.25(0.04)** | **0.25(0.03)** | **0.24(0.03)** |
| | CARS (MSR) | 3.7(0.14) | 3.7(0.13) | 3.72(0.11) | 3.71(0.11) | 3.71(0.11) | 3.71(0.13) | 3.71(0.12) | 3.73(0.11) | 3.73(0.12) | 3.74(0.1) | 3.74(0.11) | 3.75(0.11) | 3.75(0.1) | 3.75(0.09) | 3.74(0.1) | 3.72(0.33) | 3.72(0.12) | 3.71(1.01) |
