## Supporting Information 3: Table for "Benchmark of biomarker identification and prognostic modeling methods on diverse censored data"

| Known driver | LASSO | ALASSO | ENET | CB | RSF | sRSF | BHP | QV | CARS (MED) | CARS (MSR) |
|---|---|---|---|---|---|---|---|---|---|---|
| ATG5 | 0 | 0 | 0 | 0 | 0 | 0 | 0 | 0 | 0 | 0 |
| DLEU1 | 9 | 7 | 8 | 10 | 1 | 3 | 9 | 10 | 0 | 0 |
| FGF14 | 0 | 0 | 0 | 0 | 1 | 0 | 0 | 0 | 0 | 0 |
| FGF22 | 0 | 0 | 0 | 0 | 1 | 0 | 0 | 0 | 0 | 0 |
| FGF5 | 0 | 0 | 0 | 0 | 0 | 0 | 0 | 0 | 0 | 0 |
| FGFRL1 | 0 | 0 | 0 | 0 | 0 | 0 | 0 | 0 | 0 | 0 |
| FOXA3 | 0 | 0 | 0 | 0 | 0 | 0 | 0 | 0 | 0 | 0 |
| FOXF1 | 0 | 0 | 0 | 0 | 2 | 0 | 0 | 0 | 0 | 0 |
| FOXF2 | 0 | 0 | 0 | 0 | 0 | 0 | 0 | 0 | 0 | 0 |
| FOXG1 | 0 | 0 | 0 | 0 | 1 | 0 | 0 | 0 | 0 | 0 |
| FOXI1 | 1 | 0 | 1 | 0 | 4 | 0 | 0 | 0 | 0 | 0 |
| FOXK2 | 0 | 0 | 0 | 0 | 3 | 0 | 0 | 0 | 0 | 0 |
| FOXL2 | 0 | 0 | 0 | 0 | 0 | 0 | 0 | 0 | 0 | 0 |
| FOXN4 | 0 | 0 | 0 | 0 | 2 | 0 | 0 | 0 | 0 | 0 |
| FOXP3 | 0 | 0 | 0 | 0 | 8 | 1 | 0 | 0 | 0 | 0 |
| FOXR1 | 0 | 0 | 0 | 0 | 0 | 0 | 0 | 0 | 0 | 0 |
| IGF1R | 0 | 0 | 0 | 0 | 1 | 0 | 0 | 0 | 0 | 0 |
| IGF2BP1 | 0 | 0 | 0 | 0 | 0 | 0 | 0 | 0 | 0 | 0 |
| MMP19 | 2 | 0 | 1 | 0 | 3 | 0 | 0 | 0 | 0 | 0 |
| MMP20 | 0 | 0 | 0 | 0 | 4 | 0 | 0 | 0 | 0 | 0 |
| MMP27 | 0 | 0 | 0 | 0 | 10 | 5 | 0 | 0 | 3 | 0 |
| MMP8 | 0 | 0 | 0 | 0 | 2 | 2 | 0 | 0 | 1 | 0 |
| PLA2G3 | 0 | 0 | 0 | 0 | 0 | 0 | 0 | 0 | 0 | 0 |
| PLA2G4A | 0 | 0 | 0 | 0 | 2 | 0 | 0 | 0 | 0 | 0 |
| PLA2G4D | 0 | 0 | 0 | 0 | 0 | 0 | 0 | 0 | 0 | 0 |
| PSMD13 | 0 | 0 | 0 | 0 | 0 | 0 | 0 | 0 | 0 | 0 |
| PSMD4 | 0 | 0 | 0 | 0 | 1 | 0 | 0 | 0 | 0 | 0 |
| SQLE | 0 | 0 | 0 | 0 | 2 | 1 | 0 | 0 | 0 | 0 |
| TOP1 | 0 | 0 | 0 | 0 | 0 | 0 | 0 | 0 | 0 | 0 |
| TOP3B | 2 | 1 | 1 | 3 | 10 | 10 | 10 | 10 | 0 | 0 |
