## Supporting Information 1: Appendix for "Benchmark of biomarker identification and prognostic modeling methods on diverse censored data"

### Supporting information for "Benchmark of biomarker identification and prognostic modeling methods on diverse censored data"

Wesley Fletcher* and Samiran Sinha

Department of Statistics, Texas A&M University, College Station, Texas, United States of America

* Corresponding author
 (WF)

### Additional studies

We have performed additional studies to address potential concerns with our study designs. Corresponding R code for performing these studies can be found in our repository within the "supplementary" folder.

### Impact of RSF tuning parameters on performance

The RSF has two main hyperparameters that require tuning, namely the minimum node size and number of candidate features for splitting each node upon. In all components of our benchmark study, we used values of 8 and 403 respectively for these parameters, which were decided upon by an additional brief simulation study described in the main manuscript following the definition of the RSF. To quantify how this choice may have affected the method's performance, we performed an additional simulation study.

#### Method

We generated 20 synthetic datasets using the same procedure in our first setting of simulations described in the main manuscript (1000 features, 200 observations for model training, 100 observations for evaluating out-of-sample prediction) with data characteristics $s = 0.05$, $\alpha = 0.5$, and $\gamma = 1$. Each dataset was used to fit three RSF models, each using one assignment of tuning parameters: The `randomForestSRC` package default hyperparameters (32 candidate split features, minimum node size 15), our *a priori* tuned hyperparameters used in the main study, or hyperparameters tuned at runtime to maximize out-of-sample concordance on the individual training dataset using a search algorithm provided in the `randomForestSRC` package. Once fit, each RSF was evaluated using out-of-sample CI and computation time (in the third case, this includes the time required to perform the search for best hyperparameters). Summary statistics of these metrics—formatted as mean(standard deviation)—for the three hyperparameter cases are provided in Table 9.

**Table 9. CI and computation time for three settings of RSF hyperparameters.**

| Hyperparameters | CI | Computation time (in seconds) |
|---|---|---|
| Default | 0.52(0.03) | 83.24(0) |
| *a priori* | 0.51(0.03) | 100.8(8.49) |
| Tuned at runtime | 0.51(0.03) | 1191.47(125.73) |

Better performance is shown by having larger CI and shorter computation time. CI: concordance index.

#### Results

It can be seen in Table 9 that there is very little variation in predictive ability (CI) between the settings of hyperparameters, but a tenfold increase in computation time when hyperparameters are tuned for each individual dataset at runtime.

### Comparing held-out and out-of-bag predictive metrics in RSF

The out-of-sample predictive performance of the RSF can be done in two similar ways, either by evaluating using out-of-bag (OOB) or out-of-sample (held-out) data. OOB data is the observations withheld from fitting a single tree in the ensemble while held-out data is data that is not used in fitting any trees. For consistency with the other methods, we evaluated predictive performance using held-out data in all portions of the main study, but evaluating with OOB data is a common practice in real applications. We have quantified the difference between OOB and held-out predictive evaluation in a simulation study.

#### Method

We generated 20 synthetic datasets in the same way as in our first setting of simulations in the main manuscript using 1000 features, 200 in-sample observations for fitting an RSF model, 100 held-out observations for evaluating held-out predictive ability of the resulting model, data characteristics $s = 0.05$, $\alpha = 0.5$, and $\gamma = 1$. Each RSF was fit using our *a priori* hyperparameters (minimum node size of 8, and 403 candidate splitting features at each node). Evaluation was done with CI in two ways: Firstly, evaluating each in-sample observation's concordance or discordance using only the trees in the forest for which the observation is OOB, and secondly using the held-out data in the full ensemble. Table 10 contains sample quantiles and other summary statistics for the distributions of metric values for both OOB and held-out evaluations of CI on the 20 replications.

**Table 10. Distributions of OOB and held-out CI for the RSF method.**

| Metric | Min. | 1st Quartile | Median | Mean | 3rd Quartile | Max. |
|---|---|---|---|---|---|---|
| OOB | 0.44 | 0.49 | 0.51 | 0.51 | 0.52 | 0.57 |
| Held-out | 0.46 | 0.49 | 0.51 | 0.51 | 0.53 | 0.57 |

OOB: out-of-bag, CI: concordance index, RSF: random survival forest.

#### Results

We can see in Table 10 that the distributions of CI values are remarkably similar between evaluating using OOB and held-out observations. This suggests that the discrepancy between using one means of evaluation over the other is small.

### Effect of PFS on embedded methods' performances in the real data analysis

We chose to use PFS in the second setting of simulations and real data analysis to reduce the data dimensionality of the BLCA cohort to be more similar to that of our first setting of simulations. To investigate how this choice may have affected the downstream modeling performance of the examined methods, we have performed an additional study comparing the embedded methods' performances on both the 20,240 mRNA features remaining after data cleaning and the size 3,000 feature subset retained by PFS to quantify the procedure's impact on prognostic modeling.

#### Method

To quantify the downstream effect of PFS on prognostic modeling, we randomly partitioned the 423 observations of the BLCA cohort into 10 mutually exclusive folds, each containing about 42. The embedded methods were performed 10 times on both the full 20,240 feature and reduced 3,000 feature datasets of the BLCA cohort, each time using 9 of the 10 folds for training and the remaining fold for evaluating out-of-sample CI. Computation time was also recorded while performing the methods. Boxplots of these metrics for all methods on the data both before and after PFS are shown in Fig 10.

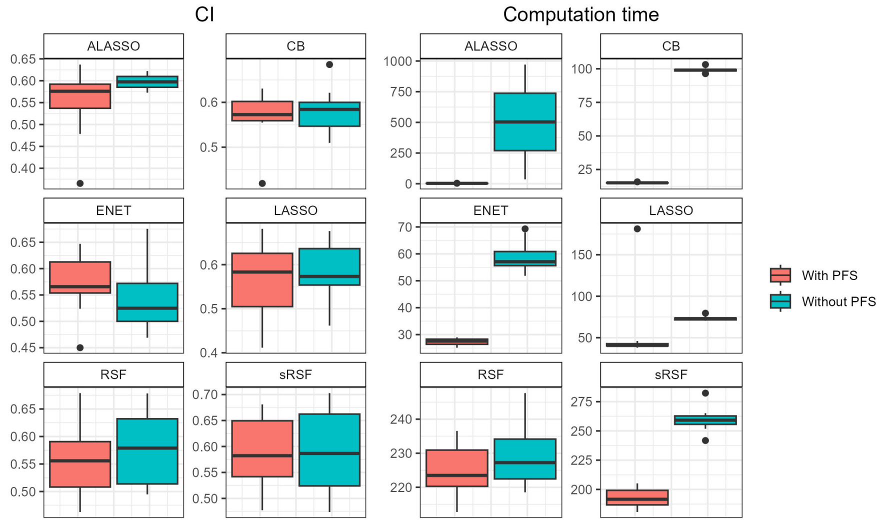


**Fig 10. Embedded method CI and computation time on BLCA cohort with and without PFS.** (Left) CI. (Right) Computation time. Plots are faceted by examined method. CI: Concordance index. Computation time was measured in seconds.

#### Results

The boxplots in Fig 10 of out-of-sample CI and computation time of the methods on the data before and after PFS show that most of the embedded methods experience a slight decay in predictive ability and a substantially large decrease in computation time when PFS is done prior to method execution. Most notably, the ALASSO, ENET, and CoxBoost (CB) methods take about twice as long or more to compute when all 20,240 features are used (no PFS) versus when only the 3,000 features retained by PFS are used. The RSF-based methods had similar computation times for both the no-PFS and with-PFS cases.

#### Retention of true signals by PFS under varying signal-to-noise ratios

Another similar question of interest is the relationship between the effect of a true signal (which $|\beta_i|$ is proportional) and its rate of retention by our PFS step (the CARS filter). We investigated this question with an additional simulation study.

##### Method

We considered all 432 observations and 20,240 mRNA features from the cleaned BLCA cohort with corresponding true observed event times and censoring indicators. Call the covariate matrix $X$. We performed CoxBoost on the real data, and found 59 nonzero regression coefficients. We used ten times of the estimated coefficients as the ground truth and generated twenty synthetic datasets for a brief simulation study. The nonzero elements of $\beta$ had absolute values between 0.15 and 1.3. We performed the following procedure 20 times: Synthetic survival times $T$ were generated for each observation following the Weibull distribution with shape 2 and scale $300 \exp(-X^\top \beta/2)$, similarly to the second setting of simulations in the main manuscript. Individual censorship times $C$ were generated to follow $\text{Unif}(Q(0.5), Q(0.9))$ where $Q(r)$ is the $r$-th sample quantile of $T$. Observed event times were $Y = \min(T, C)$ and censoring indicators were $\Delta = \mathbb{I}(T < C)$. We performed PFS on $(X, Y, \Delta)$ and recorded which of the 59 true signals were not removed by the procedure. Fig 11 shows the relationship between signal-to-noise ratio ($|\beta|$) and retention rate of the 59 features over the 20 repetitions of the study.

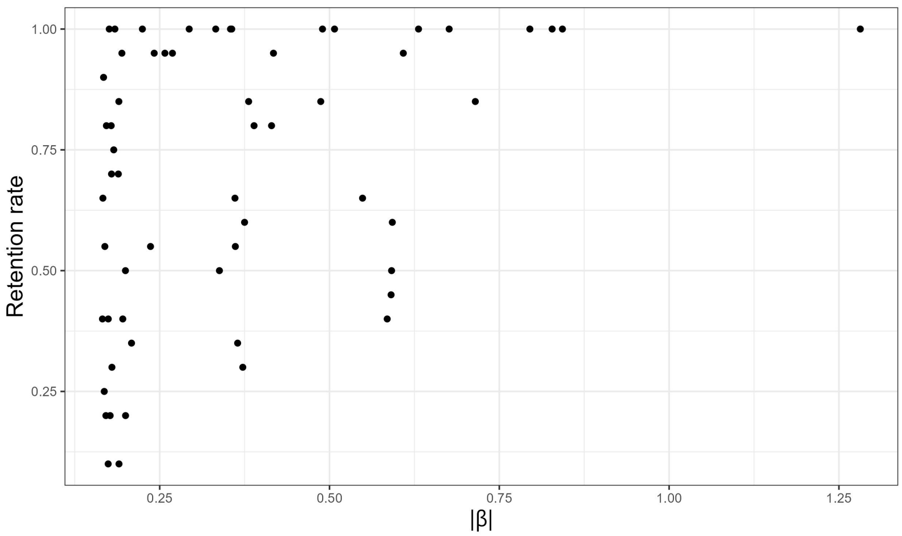


**Fig 11. Retention rate by PFS versus signal-to-noise ratio of the true signals.** Each individual point of the scatterplot corresponds to one of the 59 features, and shows the relationship between its signal-to-noise ratio (for which $|\beta|$ is a proxy) and its rate of retention by PFS in the 20 performed replications.

#### Results

The scatterplot in Fig 11 shows a general increase to retention rate as the signal strength increases. Interestingly, many of the features with lesser signals still have high retention rates by PFS. As CARS prioritizes the selection of features with greater marginal correlation with the outcome and lesser multi-colinearity, the features with higher retention rates in spite of their lesser signals likely have very little correlation with other features.

### Sensitivity analysis of PFS

In the real data analysis, we chose to perform PFS once using all observations instead of per-fold so that feature selection performance was done on identical feature sets every fold. This may have introduced some downstream data leakage, as the feature set was determined using observations that will be used to evaluate method performance. We examined how much leakage may have occurred by evaluating the stability of PFS when done per-fold.

#### Method

We used a subset of 19,805 mRNAs features $X$ from the BLCA cohort after removing very low varying features (fewer than 11 unique values), observed survival times $Y$, and censoring indicators $\Delta$. The 423 observations were placed into 10 similarly sized folds using the same assignments from the real data analysis. The 485 features that were removed had between 1 and 10 unique values inclusive, and were removed so that no features were constant among any selection of 9/10 folds. PFS was performed 10 times on $(X, Y, \Delta)$, where each time 1 fold of observations was excluded and the remaining 9 were standardized so that each feature had mean 0 variance 1. Let $S_i$ denote the set of selected features from PFS after excluding the $i$-th fold of observations, $S_0$ be the set of features kept by PFS when performed on all standardized observations (the same set used in the real data analysis and setting-II of the simulation studies), and $S_0' \subset S_0$ be the set of important features identified from existing literature, detailed in Table 2. Note that all $\#(S_i) = \#(S_0) = 3{,}000$ and $\#(S_0') = 30$, where $\#(A)$ is the cardinality of $A$. We recorded the values of $\#(S_i \cap S_0)/3{,}000$ for $i = 1, \cdots, 10$ and $\#(S_i \cap S_j)/3{,}000$ for all $i, j \in \{1, \cdots, 10\}$, which are equivalent to Dice index between the respective sets. We also recorded $\#(S_i \cap S_0')$ for $i \in \{1, \cdots, 10\}$. Summaries of these values are presented in the Table 11.

**Table 11. Distributions of Dice coefficients and a similar metric for the PFS step.**

| Metric | Min. | 1st Quartile | Median | Mean | 3rd Quartile | Max. |
|---|---|---|---|---|---|---|
| $\#(S_i \cap S_0)/3{,}000$ | 0.6207 | 0.7198 | 0.8303 | 0.7770 | 0.8443 | 0.8610 |
| $\#(S_i \cap S_j)/3{,}000$ | 0.4673 | 0.6107 | 0.6830 | 0.6821 | 0.7843 | 0.8160 |
| $\#(S_i \cap S_0')$ | 19 | 22 | 26 | 25 | 27.75 | 29 |

The first two metrics are equivalent to Dice coefficient between the corresponding sets, which ranges from 0 (no set overlap) to 1 (identical feature sets). The third metric ranges from 0 (no important features kept) to 30 (all important features kept). PFS: preliminary feature selection.

#### Results

We can see in Table 11 that a majority of the $S_i$ pairs have at least 3/5 overlap with each other, 4/5 overlap with $S_0$, and include 5/6 of the important features $S_0'$. This suggests that PFS may be stable and that there may not be much downstream affect from performing it on the full data versus per-fold.
